# *Arabidopsis* cysteine protease RD19 limits RBOHD-generated ROS production in pattern triggered immunity

**DOI:** 10.64898/2026.10.02.755912

**Authors:** Giuliana Hessler, Melissa Mantz, Federica Locci, Jaqueline Bautor, Pitter F. Huesgen, Jane E. Parker

## Abstract

Plants perceive pathogenic threats through receptors localized at the plasma membrane and inside cells to initiate pattern-triggered immunity (PTI) and effector-triggered immunity (ETI), respectively. PTI and ETI pathways are interconnected and partially converge on the EDS1-PAD4-ADR1 branch which stimulates anti-pathogen defence. Besides these canonical immune pathways, proteolysis plays a pivotal role in immunity by regulating the abundance and activities of diverse components. We previously identified *Arabidopsis* papain-like cysteine protease RD19 as a PAD4 interactor and showed that RD19 promotes resistance to host-adapted pathogens. Here, we investigated the contribution of RD19 to bacterial flg22 and nlp20 elicited PTI responses which exhibit, respectively, partial and high *PAD4* dependence. We show that although PAD4 and RD19 both promote PTI, they have separate functions. In resting (non-triggered) and nlp20-triggered tissues, RD19 limits RBOHD protein accumulation and consequently ROS production. Using N-terminomic enrichment and protease cleavage site identification methods we identify candidate RD19 substrates in resting and nlp20-responding tissues. Our analysis shows that two immune-related aspartyl proteases AED3 and AED2/ASPG1 are likely direct RD19 targets. This study reveals a role of RD19 in ROS regulation and advances our understanding of proteolysis and dynamic RD19 substrate targeting in plant immunity.

**Highlight:** *Arabidopsis* RD19 cysteine protease modulates accumulation of the immune signalling hub protein RBOHD and its generation of ROS. RD19 processes multiple immune-related proteins and promotes defence potentiation in pattern-triggered immunity.

## Introduction

Plants have evolved a two-layered immune system to combat pathogen infection. The first layer, called pattern triggered immunity (PTI), is activated when plasma membrane receptors (PRRs) detect pathogen- or damage-associated molecular patterns (PAMPs or DAMPs) (Dangl and Jones, 2001). PRRs include receptor kinases (RKs), such as *Arabidopsis* FLS2 which recognizes the bacterial flagellin epitope flg22, and receptor-like proteins (RLPs), such as RLP23 which detects necrosis- and ethylene-inducing peptide 1–like proteins (NLPs) nlp20 and nlp24 (Gómez-Gómez and Boller, 2000; Chinchilla *et al*., 2007; Böhm *et al*., 2014; Albert *et al*., 2015). Both receptors form an induced signalling complex with the RK co-receptor BRI1-ASSOCIATED RECEPTOR KINASE 1 (BAK1) while RLP23 constitutively associates with RK SUPPRESSOR OF BIR1 1 (SOBIR1), providing a crucial intracellular kinase signalling domain (Chinchilla *et al*., 2007; Albert *et al*., 2015). In *Arabidopsis thaliana* (hereafter *Arabidopsis*), PTI can be broadly separated into early- (within minutes) and late-phase (hours to days) responses. Early defence signalling processes include the generation of apoplastic reactive oxygen species (ROS) mainly mediated by NADPH/RESPIRATORY BURST OXIDASE PROTEIN D (RBOHD) (Kadota *et al*., 2015) general stress induced transcriptional reprogramming (Jacob *et al*., 2018; Bjornson *et al*., 2021), transient Ca^2+^-influxes into cells and phosphorylation of MITOGEN-ACTIVATED PROTEIN KINASES (MAPKs) (DeFalco and Zipfel, 2021). PTI late-phase responses include prolonged defence gene expression changes and induced resistance to subsequent pathogen infection (Zipfel *et al*., 2004; Bjornson *et al*., 2021; Pruitt *et al*., 2021). The second immunity layer is activated by pathogen virulence factors (effectors) that are delivered into host cells to dampen PTI and promote infection (Dangl and Jones, 2001; Ngou *et al*., 2022; Jones *et al*., 2024). Effector-triggered immunity (ETI) is mediated mostly by intracellular nucleotide-binding leucine-rich repeat receptors (NLRs) with coiled-coil (CC) or Toll-Interleukin-1 Receptor (TIR) signalling domains (respectively, CNLs and TNLs), which recognize particular effectors or effector activities to induce a stronger immune response often culminating in localized host cell death (Dangl and Jones, 2001; Contreras *et al*., 2023; Jones *et al*., 2024).

Execution of cell death and cell-to-cell signalling require the production of ROS (Van Breusegem and Dat, 2006; Mittler *et al*., 2022; Peláez-Vico *et al*., 2022). Therefore, tight regulation of ROS production, and hence RBOHD as a regulatory hub, ensures a balanced immune response (Mittler *et al*., 2022). RBOHD is activated by Ca^2+^-dependent and Ca^2+^-independent mechanisms. For example, Ca^2+^ binds a helix-loop-helix (EF) hand motif of RBOHD and mobilizes calcium-dependent protein kinases (CPKs) such as CPK5 which phosphorylates RBOHD (Dubiella *et al*., 2013). BOTRYTIS-INDUCED KINASE (BIK1) phosphorylates RBOHD to promote its function independently of Ca^2+^ (Dubiella *et al*., 2013; Kadota *et al*., 2014; Li *et al*., 2014). As checkpoints against mis-activation, RBOHD is negatively regulated by receptor-like cytoplasmic kinases (RLCKs) as well as E3-ubiquitin ligases and proteases (Lin *et al*., 2015; Lee *et al*., 2020; Liu *et al*., 2023).

In *Arabidopsis*, ETI and PTI pathways cross-potentiate each other, with ETI serving to amplify PTI-related ROS production, Ca^2+^ and MAPK cascades and anti-pathogen defence (Ngou *et al*., 2021, 2022; Yuan *et al*., 2021). Nucleocytoplasmic heterodimers of the lipase-like proteins ENHANCED DISEASE SUSCEPTIBILITY 1 (EDS1) and PHYTOALEXIN-DEFICIENT 4 (PAD4) transduce signals from activated NLRs (Lapin *et al*., 2020) and certain PRRs, including nlp20-triggered RLP23 (Dongus and Parker, 2021; Pruitt *et al*., 2021). In this role, EDS1-PAD4 is a receptor for ribosylated nucleotide second messengers (SMs) generated by TNL and TIR-protein enzymes (Huang *et al*., 2022; Jia *et al*., 2022; Locci and Parker, 2024; Fliegmann *et al*., 2026). In both ETI and PTI, the SM-bound EDS1-PAD4 dimers function together with ACTIVATED DISEASE RESISTANCE 1 (ADR1)-family signalling (“helper”) NLRs to stimulate defence transcription which limits pathogen growth without host cell death (so-called basal immunity) (Bonardi *et al*., 2011; Castel *et al*., 2019; Lapin *et al*., 2019; Wu *et al*., 2019; Saile *et al*., 2020; Parker *et al*., 2021; Sun *et al*., 2021). Signalling via EDS1-PAD4-ADR1 is required for early and late PTI responses to nlp20 (Pruitt *et al*., 2021; Fliegmann *et al*., 2026). Also, the *Arabidopsis* EDS1-PAD4-ADR1 node operates at the borders of infection sites and contributes to propagating signals which drive systemic immunity (systemic acquired resistance (SAR)), through induced salicylic acid (SA)-dependent and SA-independent pathways (Rustérucci *et al*., 2001; Wittek *et al*., 2014; Jacob *et al*., 2023). Hence, EDS1-PAD4 represents a convergence point for cell-surface and intracellular immune receptor signalling.

Besides the canonical receptor-mediated defence pathways, additional regulators are required for precise activation and deactivation of immune components. Here, proteolysis has been proven to play an essential role (Liu *et al*., 2024). In plant immunity proteases function in various subcellular localizations including the vacuole, the nucleus and apoplastic space (Van Der Hoorn, 2008). Among the diverse range of proteases found in *Arabidopsis*, cysteine proteases, specifically papain-like cysteine proteases (PLCPs), were proposed to be key players in fine-tuning immune responses (Misas-Villamil *et al*., 2016). PLCPs belong to the MEROPS protease family C1A of clan CA that carries a conserved Cys, His and Asn catalytic triad within the substrate-binding groove of the protease domain (Rawlings *et al*., 2018). Moreover, PLCPs contain an N-terminal signal peptide (SP) for secretion and an auto-inhibitory pro-domain which requires cleavage for activation (Misas-Villamil *et al*., 2016). The role of PLCPs in immunity ranges from processing of defence-related substrates, activation of immune signalling peptides to promotion of pathogen resistance via cell death (Misas-Villamil *et al*., 2016).

*Arabidopsis* PLCP XYLEM CYSTEINE PEPTIDASE 1 (XCP1) was shown to destabilize RBOHD by degradation in the vacuole under resting state conditions (Liu *et al*., 2023). XCP1 also cleaves a caspase-like motif of PATHOGENESIS RELATED 1 (PR1) upon SA treatment, releasing a small peptide which in turn activates SA biosynthesis and systemic immunity (Chen *et al*., 2023). The activity of most cysteine proteases is regulated post-translationally by external stimuli, such as pH change, Ca^2+^-influx or oxidative stress (Fernández-Fernández *et al*., 2023). METACASPASE 4 (MC4), for example, exists in its inactive zymogen form (zMC4) in undisturbed tissues and is activated upon damage-induced Ca^2+^-influx, leading to cleavage of the endogenous peptide PROPEP1 to release the phytocytokine Pep1 (Hander *et al*., 2019). Most plant proteases studied to date self-cleave their inhibitory pro-domains under activating conditions. However, there are examples of proteases that require cleavage by other proteases for activation, for example the tomato PLCP Rcr3 which is activated by subtilase P69B (Paulus and Van der Hoorn, 2019; Paulus *et al*., 2020).

Since protease activation and substrate availability often depend on cellular or environmental stimuli, subcellular localization can be an important determinant of protease activityand tissue-or condition-specific functions. For example, XCP1 cleaves RBOHD in the vacuole but PR1 in the apoplast (Chen *et al*., 2023; Liu *et al*., 2023). PLCP RESPONSIVE TO DEHYDRATION 19A (RD19A) was shown to localize to mobile vacuole-associated vesicles and the vacuole in non-immune elicited conditions while presence of the TNL recognized *R. solanacearum* effector PopP2 (*Pseudomonas* outer protein P2) led to co-localization of RD19A and PopP2 to the nucleus (Bernoux *et al*., 2008). By contrast, a *Zea mays* RD19A-like protein localized to the root apoplast where it was targeted by a membrane-anchored *Pseudomonas* inhibitor to suppress its activity (Schulze Hüynck *et al*., 2019; Moser *et al*., 2026). These observations suggest that changes in PLCP localization contribute to their functions in different cellular contexts.

We previously identified RD19A as an interactor of *Arabidopsis* PAD4 and found that zymogen RD19A accumulated in the nucleus in the presence of PAD4 (Zeng *et al*., 2024). RD19A overexpression led to EDS1/PAD4-dependent autoimmunity, indicating that regulation of RD19A is crucial for appropriate immune activation (Zeng *et al*., 2024). Loss of RD19A and a second isoform, RD19C, compromised basal immunity to powdery mildew but not to *Pseudomonas syringae* bacteria, suggesting that RD19A and RD19C positively contribute to a subset of EDS1-PAD4-ADR1 regulated immune responses (Zeng *et al*., 2024).

Because the EDS1-PAD4 dimer is required for nlp20-activated PTI responses, and RD19A was identified as a PAD4-specific interactor, here we investigated involvement of RD19 proteases in *PAD4*-dependent and -independent PTI. We found that RD19A and RD19C limit RBOHD protein accumulation and ROS production in nlp20-triggered PTI and yet promote nlp20-triggered induced resistance to *P. syringae* bacterial and *Botrytis cinerea* fungal infections. Comparative phenotyping of *rd19ac* and *pad4* mutants established that their functions only partially overlap, reflecting likely distinctive roles in PTI. We identified RD19A amino acid cleavage preferences and integrated this information with N-terminomic profiling of RD19A substrates to identify two immune-related aspartic proteases, suggesting operation of an RD19-dependent proteolytic cascade in nlp20-triggered immunity. We propose a previously unknown role of RD19 in ROS balance and defence potentiation via newly identified substrate candidates.

## Materials and Methods

### Plant materials and growth conditions

*Arabidopsis* seeds were sown on soil fertilized with 10 mg/l Confidor WG70 (Bayer) and grown under short day conditions (10 h light, 14 h dark) at ∼150 µmol/m/s, 22 °C and 65 % humidity. The following mutant lines used in this study were previously described: *rd19ac* (Zeng *et al*., 2024), *pad4-1* (Glazebrook *et al*., 1997), *fls2* (Zipfel *et al*., 2004), *rlp23* (Pruitt *et al*., 2021), *rbohd* (Torres *et al*., 2002). Triple mutant lines *rd19ac rbohd* and *rd19ac pad4-1* were generated in this study.

### Plasmid construction and site-directed mutagenesis

In-Fusion seamless cloning (Takara Bio) was used to generate plant expression vectors. RD19A coding sequence (CDS) was amplified from Col-0 cDNA. Primers generating overhangs were designed using the Takara Bio online design tool (https://www.takarabio.com/learning-centers/cloning/primer-design-and-other-tools). The L1 MoClo vector (Moser *et al*., 2026) was used as a backbone for transient expression of RD19A and candidate substrates in *N. benthamiana*. Primers used for cloning and site-directed mutagenesis are listed in Supplementary Table S1.

### Pseudomonas syringae infection assays

Infection assays with *Pseudomonas syringae* pv. *tomato* (*Pst*) strain DC3000 were performed as previously described (Lapin *et al*., 2019), except for using bacterial cells at a density of OD_600_=0.0002. For spray inoculations with *Pst* DC3000 lacking the hormone Coronatine (*Pst* ΔCor), bacteria were diluted to a concentration of OD_600_=0.2 in 10 mM MgCl_2_ with 0.04% (v/v) Silwet-L77. Leaf disc sampling and bacteria CFU calculations were performed as previously described (Lapin *et al*., 2019) .

### Botrytis cinerea infection assays

*Botrytis cinerea* infections with strain B05.10 were performed as previously described (Prior *et al*., 2026), except using spore solutions of 5×10^5^ spores/ml and infecting at least 10 leaves per sample with two 10 µl drops of spore suspension. Lesions were imaged with the BioRad ChemiDoc™ XRS+ detection system 48 h after infection. Lesion areas were measured on photos using Fiji (Schindelin *et al*., 2012). In case of induced resistance experiments, all leaves were sprayed with elicitor solutions (0.5 µM elicitor in ddH_2_O) 24 h before infection with *B. cinerea*.

### RT-qPCR

Real time quantitative PCR was performed on cDNA using the iQ SYBR Green Supermix (BioRad). Primer sequences are described in Supplementary Table S1. RT-qPCRs were run in the CFX Connect Real-Time PCR Detection System (BioRad). The obtained Cq values were normalized to the Cq values of the housekeeping gene relative to the Col-0 mock treatment.

### MAPK phosphorylation

Assays to detect phosphorylation of MAPKs were performed as previously described (Pruitt *et al*., 2021). MAPK phosphorylation was visualized on immunoblotting using the α-p44/42 primary antibody (1:1000; Cell Signaling, #9101).

### Reactive oxygen species (ROS) measurements

ROS assays were performed as previously described (Pruitt *et al*., 2021), except using leaf discs with a diameter of 0.5 cm. Luminescence was measured over 1 h using a Centro XS^3^ LB 960 machine (Berthold Technologies).

### Protein expression analyses in *Arabidopsis* and *Nicotiana benthamiana*

For protein expression analysis, four 1 cm leaf discs of transiently transformed *Nicotiana benthamiana* or 2 leaves of *Arabidopsis* were harvested in Eppendorf tubes containing metal beads and immediately frozen in liquid nitrogen. A Qiagen TissueLyser II was used at 30 Hz for 1 min to grind plant material. Samples were resuspended in 2x Laemmli buffer (60 mM TRIS pH 6.8, 4 % (w/v) SDS, 200 mM DTT, 20 % (w/v) glycerol, 0.2 % (w/v) bromophenol blue), boiled at 95 °C for 10 min and centrifuged at 13,300 rpm for 1 min. Supernatants were collected and protein concentrations determined by Pierce 660 nm assay (ThermoFisher). Each sample was diluted with 2x Laemmli buffer to a protein concentration of 60 µg and separated by SDS-PAGE. RBOHD protein was detected on immunoblots using α-RBOHD (1:5000; Agrisera, AS152962), followed by α-rabbit-HRP secondary antibody (1:5000; Cell Signalling, 70745). Expression of RD19A-3xHA (WT and catalytically inactive mutant) was detected by α-HA (1:5000; Roche, 11867423001) and α-rat-HRP (1:5000; Sigma-Aldrich, AP136P) antibodies and RD19 candidate substrates were detected using α-GFP (1:5000, Roche, 11814460001) and α-mouse-HRP (1:5000, Cell Signalling, 70765). Equal protein loading was monitored by Ponceau staining.

### Stomata aperture index (SAI) measurements

Guard cell aperture assays were performed as previously described (Eisele *et al*., 2016). Briefly, epidermal peels were obtained from 4-5-week-old *Arabidopsis* plants by peeling the abaxial side of rosette leaves using laboratory tape. Epidermal peels were placed in MES/KOH buffer (5 mM KCl, 50 µM CaCl_2_, 10 mM MES pH 6.15) for 2 h and then transferred to fresh MES/KOH solutions containing 1 µM of the elicitors flg22 or nlp20, or ddH_2_O for mock treatments for 2 h. Peels were then stained and fixed in 4 % Paraformaldehyde (PFA) solutions with 1 µM rhodamine 6G. Epidermal peels were imaged using a Zeiss LSM780 confocal microscope. Rhodamine 6G was imaged with λ_Ex_ = 514 nm, λ_Em_ = 535-580 nm. Fiji software (Schindelin *et al*., 2012) was used to measure length and width of guard cells and the stomatal aperture index (SAI, width/length) was calculated.

### Proteomic Identification of Protease Cleavage Sites (PICS)

*Agrobacterium tumefaciens* transformed with 35S::RD19A^WT^-3xHA and 35S::RD19A^C156A_C159A^-3xHA were transiently expressed in *N. benthamiana* at OD_600_ = 1 for 48h. Leaves were placed in beakers containing ddH_2_O and apoplastic fluids (AFs) were isolated by applying a vacuum of 60 mbar for 5 min and centrifuging infiltrated leaves at 800 x g for 15 min to collect AFs. The PICS experiment testing the activity of RD19A^WT^ against *E. coli* proteome-derived peptide libraries, using the catalytically inactive and an E64-pretreated inactive RD19A^WT^ as controls, was then performed as described (Demir *et al*., 2022*a*). See supplementary information for experimental details and mass spectrometry data acquisition. Peptides were identified with MaxQuant (version 2.4.2.0) (Tyanova *et al*., 2016), with library-specific settings as described (Demir *et al*., 2022*a*). Cleavage windows were reconstructed from the observed peptides only identified after incubation with active RD19A, but none of the controls using the perl script PINCIS (version 12, downloaded from https://pincis.sourceforge.io) (Demir *et al*., 2022*a*) and sequence specificity visualized as iceLogos (Colaert *et al*., 2009).

### High-efficiency Undecanal-based N-Termini EnRichment (HUNTER)

For HUNTER experiments, 4–5-week-old *Arabidopsis* Col-0 and *rd19ac* leaves were infiltrated with nlp20 solutions at 1 μM and snap frozen and ground after 10 min or 3 h of treatment. Untreated controls were sampled immediately. Ground frozen leaf material was suspended in denaturing protein extraction buffer (6M guanidine, 100 mM HEPES, pH 7.5, supplemented with Halt Protease-Inhibitor-Cocktail, Thermo) and processed essentially as described (Demir *et al*., 2022*b*). See supplementary information for details on sample processing, mass spectrometry data acquisition and peptide identification.

Peptides identified in HUNTER experiments were further analysed with in-house R-Scripts written and executed with RStudio (Racine, 2012). Peptide sequences were matched to the corresponding protein models downloaded from the UniProt database, identifiers and protein functional annotation retrieved and start and end positions within the protein sequence annotated and the cleavage window (AAs before and after the cleavage) reconstructed. Subcellular localization and the presence of targeting sequences was predicted with TargetP2.0 (Almagro Armenteros *et al*., 2019) . Redundant peptide identifications from different charge states and different Met oxidation were aggregated. The dataset was then limited to N-terminally blocked, semi-tryptic peptides reliably quantified in at least three replicates of each condition were retained for quantitative analysis. In addition, we considered peptides only observed in one genotype, if observed in least 3 replicates at one timepoint. For comparative statistics, linear models with empirical Bayes moderation were applied to calculate moderated t-statistics and p-values for each peptide with the R-Package limma (Linear Models for Microarray Data) (Ritchie *et al*., 2015) as provided in the Bioconductor package version 3.20 (Robinson *et al*., 2010). Contrasts were defined to compare wild-type and mutant samples separately for each time point. Peptides were considered significantly altered in abundance with an increase of at least 50% in one condition relative to the other supported by a limma-moderated p-value < 0.05. Volcano plots were generated to visualize differences in peptide abundance. The Venn diagram was generated using InteractiVenn (Heberle *et al*., 2015).

### Analysis of publicly available RNAseq and single-cell RNAseq datasets

Publicly available RNAseq datasets used for gene expression analysis in this study were downloaded from the Sequence Read Archive (SRA) of NCBI. Expression data of nlp20-treated plants was retrieved from SRP201971 (Wan *et al*., 2019) and flg22-treated plants from SRP082532 (Birkenbihl *et al*., 2017). A single cell RNAseq dataset of *Arabidopsis* Col-0 infected with *Colletotrichum higginsianum* was used to examine *RD19A* and *RD19C* gene expression (Tang *et al*., 2023). Data were downloaded from the ENA website under the project ID PRJEB61052.

### AlphaFold3 structural modelling

Structures of AED3 and ASPG1 were predicted in their full length and cleaved forms using the AlphaFold3 Server (Abramson *et al*., 2024). Predicted structures were downloaded and the software ChimeraX was used for visualization and superimposition (Pettersen *et al*., 2004).

## Results

### RD19 regulates nlp20-triggered ROS production

*Arabidopsis* RD19A was previously identified as a PAD4 interactor and analysis of a *rd19a rd19c* double mutant in accession Col-0 revealed a positive contribution of isoforms RD19A and RD19C to some but not all *PAD4*-dependent ETI and basal immune responses (Zeng *et al*., 2024). Here, we aimed to test whether RD19 and PAD4 functionally align in PTI, in which PAD4 has a major role in promoting early and late defences to the PAMP nlp20 and a positive contribution to stomatal responses induced by PAMP flg22 (Pruitt *et al*., 2021; Wang *et al*., 2024; Fliegmann *et al*., 2026). As observed for ETI (Zeng *et al*., 2024), of the four isoforms RD19A, B, C and D, only expression of *RD19A* and *RD19C* increased upon nlp20 treatment of leaves, whereas flg22 treatment marginally increased *RD19A* and decreased *RD19C* expression (Supplementary Figure S1A). We therefore compared PTI responses of the *rd19a rd19c* double mutant (*rd19ac*; Zeng et al., 2024) with a newly generated triple mutant *rd19ac pad4*, using transcriptional defence reprogramming, MAPK phosphorylation and ROS production as hallmark PTI outputs.

First, mutant and wild-type Col-0 leaves were syringe-infiltrated with flg22, nlp20 or mock solution and samples harvested at 3 hpi for gene expression analysis. Col-0 *fls2* and *rlp23* null mutant lines were tested alongside as non-responsive receptor controls for flg22 and nlp20 treatments, respectively. Flg22 induced expression of *FLG22-INDUCED RECEPTOR-LIKE KINASE 1* (*FRK1*) to a similar level in *pad4*, *rd19ac*, *rd19ac pad4* and Col-0 (Figure 1A). *PHYTOALEXIN DEFICIENT 3* (*PAD3*), encoding an enzyme required for phytoalexin biosynthesis, was used as a marker for nlp20-triggered transcriptional defence (Pruitt *et al*., 2021). While Col-0 and *rd19ac* plants showed similarly induced expression of *PAD3*, the *pad4* and *rd19ac pad4* mutants failed to induce *PAD3* expression (Figure 1A). Therefore, while *PAD4* and *RD19A, C* are dispensable for flg22 immune marker gene induction their activities diverge in response to nlp20.

**Figure 1.**
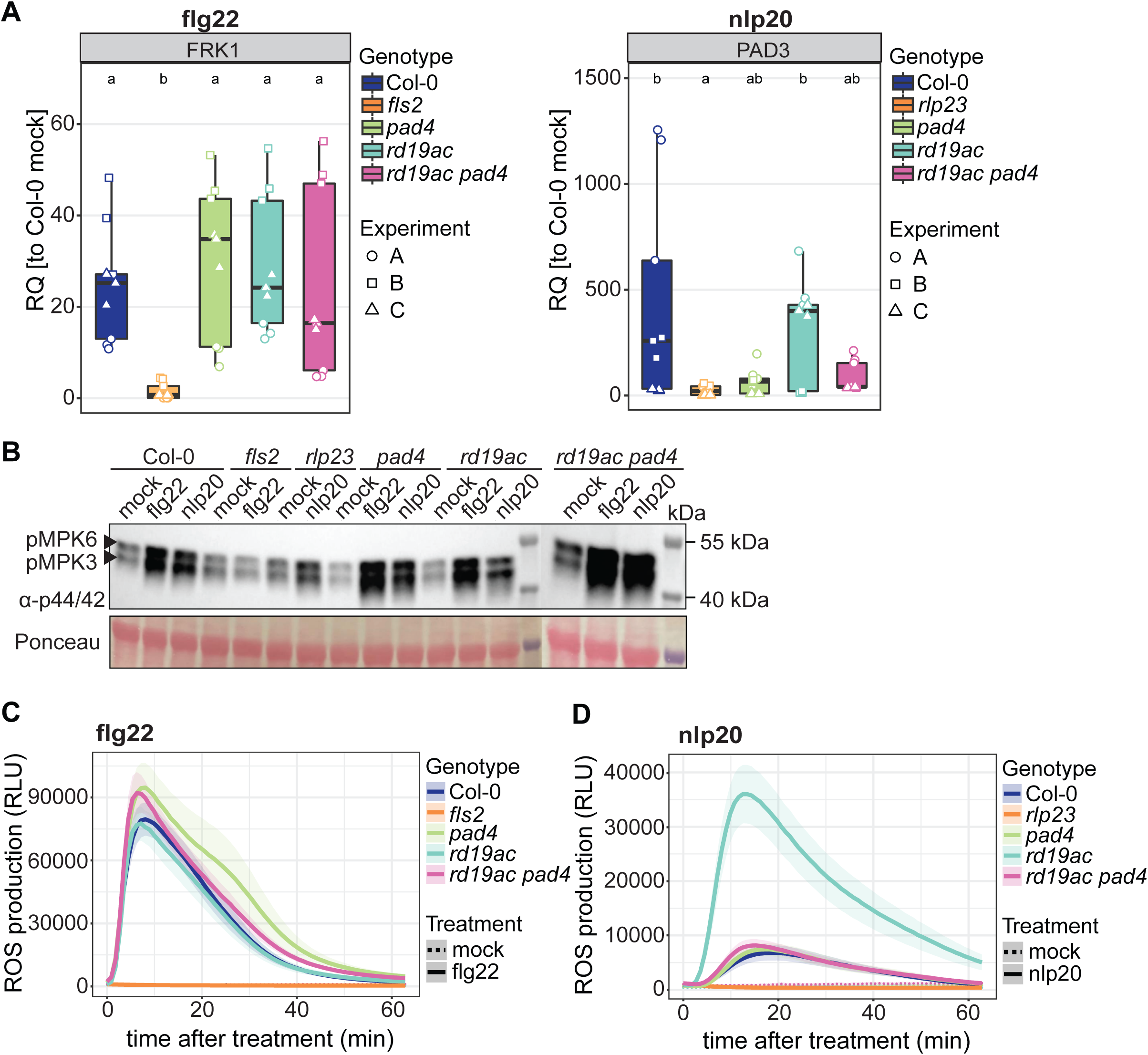
RD19 regulates the ROS burst but not defence marker gene expression or MAPK activation in response to nlp20. **A)** Expression of *FRK1* and *PAD3* 3 h after flg22 and nlp20-treatment, respectively, determined by RT-qPCR in 4–5-week-old plants. Relative quantification is represented in boxplots with dots of different shapes indicating biological replicates. Data were normalized to Col-0 mock conditions in each biological replicate. *UBI5* was used as a reference gene. Genotypes sharing letters above boxplots do not show statistically significant differences (Kruskal-Wallis and Nemenyi-Test, p < 0.05, n=9). Centre lines of boxplots indicate the median, outer edges the 25^th^ and 75^th^ percentiles and whiskers 1.5 x IQR. **B)** MAPK activation was assessed by immunoblotting with phospho-specific p44/42 antibody. Samples were taken 15 min after infiltration. A Pierce assay was performed to load equal amounts of protein and Ponceau-stained Rubisco served as a loading control. Similar MAPK activation was observed in three independent biological replicates. **C)** and **D)** ROS production measured over 60 min in leaf discs treated with mock or elicitor solutions. Shown are mean relative light units after **C)** flg22 and **D)** nlp20 treatment with standard error distributions around curves. n=24 (flg22 and nlp20) and n=6 (mock) leaf discs from three independent biological replicates.

We next tested MAPK phosphorylation in Col-0, *fls2*, *rlp23*, *pad4*, *rd19ac* and *rd19ac pad4* leaves as an early PTI output (DeFalco and Zipfel, 2021; Bender and Zipfel, 2023). MAPK phosphorylation was analysed by an immunoblot assay at 15 min after treatment with flg22 or nlp20 (Figure 1B). Both treatments induced phosphorylation of MAPK in Col-0, while the *fls2* and *rlp23* mutants exhibited weaker MPK3 and MPK6 activation, comparable to mock treatments (Figure 1B). As previously reported (Pruitt *et al*., 2021; Fliegmann *et al*., 2026), *pad4* did not diminish MAPK phosphorylation after either elicitor treatment. We found *rd19ac* and *rd19ac pad4* mutants to be similarly unaffected (Figure 1B). Therefore, both PAD4 and RD19A/C are dispensable for flg22- and nlp20-mediated MAPK phosphorylation.

We tested the production of ROS upon flg22 and nlp20 treatments in leaves of Col-0 and the above mutants as a further early PTI output. Flg22 treatment induced ROS production with a peak at 10 min in Col-0, *pad4*, *rd19ac*, and *rd19ac pad4*, but not in the *fls2* mutant (Figure 1C). A comparable ROS burst was reflected by statistical analysis performed on the area under the curve (AUC) for the tested lines (Supplementary Figure S1B). Nlp20 treatment induces a generally weaker ROS burst compared to flg22 in Col-0 plants (Pruitt *et al*., 2021). We found this was also the case but, contrasting to previous reports (Pruitt *et al*., 2021; Fliegmann *et al*., 2026), *pad4* did not display reduced ROS production compared to Col-0 upon nlp20 treatment, whereas no ROS was produced in *rlp23* under our conditions (Figure 1D; Supplementary Figure S1C). Notably, *rd19ac* generated a significantly higher ROS burst compared to Col-0 in response to nlp20 (Figure 1D). This amplified ROS response was dependent on *PAD4* as indicated by its loss in *rd19ac pad4* (Figure 1D). Statistical significance of increased ROS production in *rd19ac* was verified by AUC measurements (Supplementary Figure S1C). Taken together, these results suggest that RD19 and PAD4 fulfil different roles in PTI, with RD19A and RD19C potentially acting as a negative regulator of nlp20-induced ROS.

### RD19 limits RBOHD protein accumulation

The NADPH-oxidase enzyme RBOHD is a major generator of apoplastic ROS in plant responses to biotic stress (Torres *et al*., 2002; Torres and Dangl, 2005; Nühse *et al*., 2007; Qi *et al*., 2017). We therefore tested whether the observed increased ROS burst in *rd19ac* plants upon nlp20 treatment may be due to altered RBOHD protein accumulation. Nlp20 treatment of Col-0 plants led to a transient increase of RBOHD protein levels at 15 min after elicitor treatment, approximating to the time of the observed ROS peak (Figure 1D; Figure 2A). By contrast, the *rd19ac* mutant showed higher basal RBOHD protein accumulation compared to Col-0 in untreated leaves (Figure 2A). A further increase in RBOHD accumulation was observed at 15 min, and this remained higher than in Col-0 at 30 min after treatment (Figure 2A). In these assays, a *rbohd* mutant control did not accumulate RBOHD protein (Figure 2A). We checked whether increased RBOHD protein in *rd19ac* was a result of higher *RBOHD* gene expression. This was not the case, since *RBOHD* expression was not affected in *rd19ac* at early (Figure 2B) and late (Supplementary Figure S2) time points following nlp20 treatment. We concluded that altered RBOHD protein accumulation in *rd19ac* is not regulated at a transcriptional level but rather due to changes in protein abundance.

**Figure 2.**
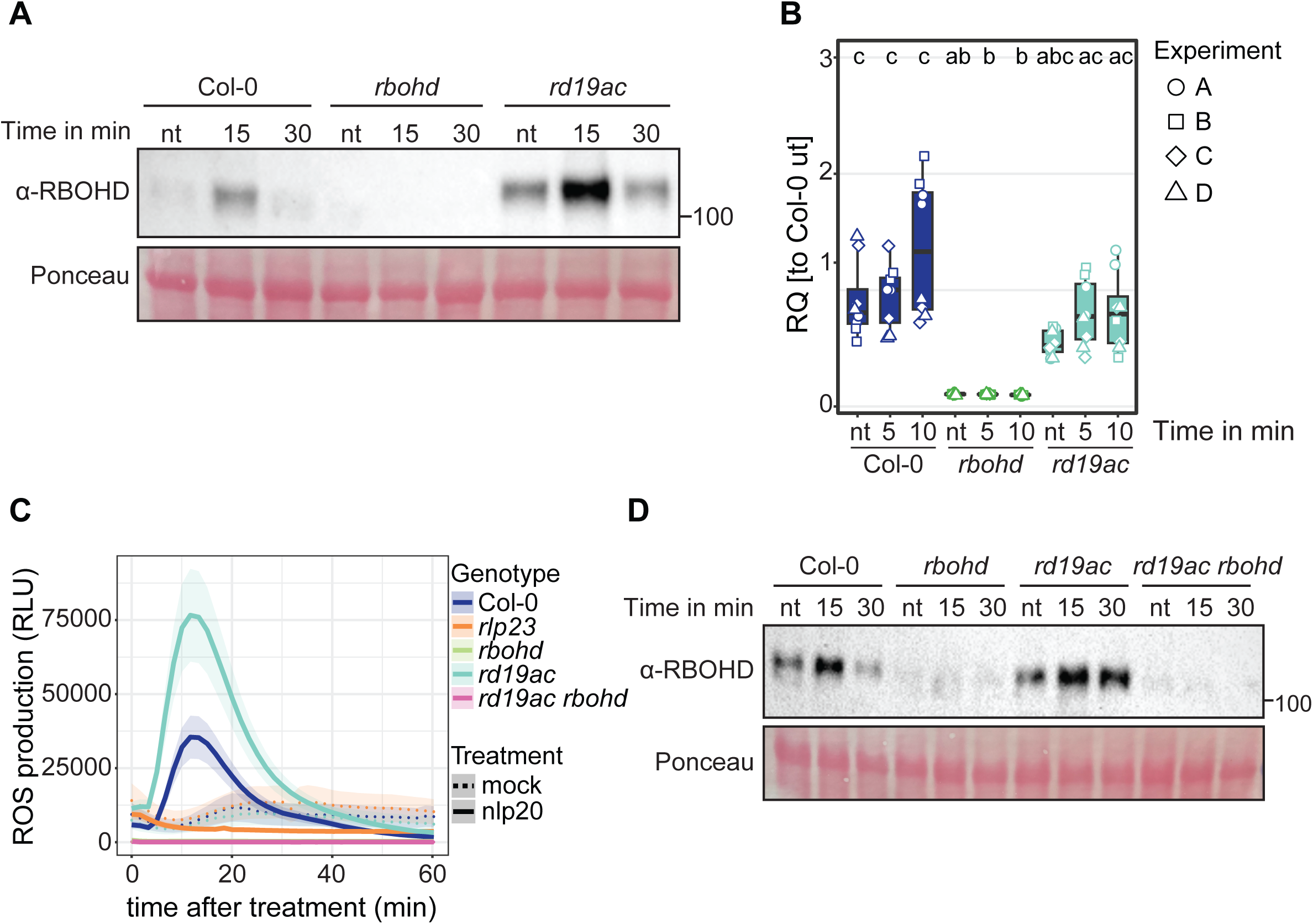
Loss of RD19A and C leads to increased RBOHD protein accumulation. **A)** RBOHD protein levels and **B)** *RBOHD* gene expression were analysed in 4.5-week-old Col-0, *rbohd* and *rd19ac* leaf extracts. Centre lines of boxplots indicate the median, outer edges the 25^th^ and 75^th^ percentiles and whiskers 1.5 x IQR. **A)** Protein was detected by immunoblotting with α-RBOHD antibody in non-treated (nt) tissue, or 15 min and 30 min after nlp20 infiltration. A Pierce assay was performed to load equal amounts of protein and Ponceau-stained Rubisco served as a loading control. Similar RBOHD protein levels were observed in four independent biological replicates. **B)** *RBOHD* gene expression was measured in non-treated (nt) tissue, and 5 min and 10 min after nlp20 infiltration. Relative quantification is represented in boxplots with dots of different shape indicating biological replicates. Data were normalized to Col-0 mock conditions in each biological replicate. *UBI5* was used as a reference gene. Genotypes sharing letters above boxplots do not show statistically significant differences (Kruskal-Wallis and Nemenyi-Test, p < 0.05, n=8). **C)** ROS production measured over 60 min in leaf discs treated with nlp20 elicitor solution or a mock solution. Shown are mean relative light units after nlp20 treatment with standard errors shown by distribution around curves. n=24 (nlp20) and n=6 (mock) leaf discs from three independent biological replicates. **D)** RBOHD protein levels were identified by immunoblotting with RBOHD antibody in non-treated (nt) tissue, or 15 min and 30 min after nlp20 infiltration. A Pierce assay was performed to load equal amounts of protein and Ponceau-stained Rubisco served as a loading control. Similar RBOHD protein levels were observed in three independent biological replicates.

To verify whether the increased nlp20-triggered ROS burst in *rd19ac* was dependent on RBOHD, we crossed *rd19ac* with *rbohd* and tested the resulting *rd19ac rbohd* triple mutant for ROS production and RBOHD protein accumulation following nlp20 treatment (Figure 2C and Figure 2D). In both assays, the *rd19ac rbohd* mutant behaved like the *rbohd* single mutant, indicating that enhanced ROS production in nlp20-elicited *rd19ac* was mediated by RBOHD (Figure 2C and Figure 2D). We therefore concluded that RD19A and C likely restrict ROS accumulation by limiting RBOHD protein synthesis and/or turnover.

### RD19 positively regulates stomatal opening after nlp20 elicitation

Stomatal closure requires apoplastic ROS and is produced mainly by RBOHF and RBOHD (Singh *et al*., 2017; Arnaud *et al*., 2023). Since *rd19ac* mutants displayed an increased ROS burst upon nlp20 treatment and both *RD19A* and *RD19C* are highly expressed in stomatal cells following pathogen infection (Tang *et al.,* 2023; Supplementary Figure S3), we tested whether RD19A and C regulate stomatal closure in response to nlp20. Epidermal peels of Col-0, *rbohd* and *rd19ac* were placed in nlp20 elicitor or mock solution and used for analysis of stomatal function by measuring the stomata aperture index (SAI). In Col-0 plants, nlp20 induced a significant SAI reduction, representing increased closure of stomata, while *rbohd* mutants displayed a SAI similar to mock treatment and therefore a full deficiency in stomatal closure (Figure 3A). While the SAI in *rd19ac* mutants was lower in response to nlp20 compared to mock treatment, (Figure 1D and Figure 3A), the elicitor effect was weaker than for Col-0 plants, as stomata in *rd19ac* exhibited enhanced closure under mock conditions (Figure 3A).

**Figure 3.**
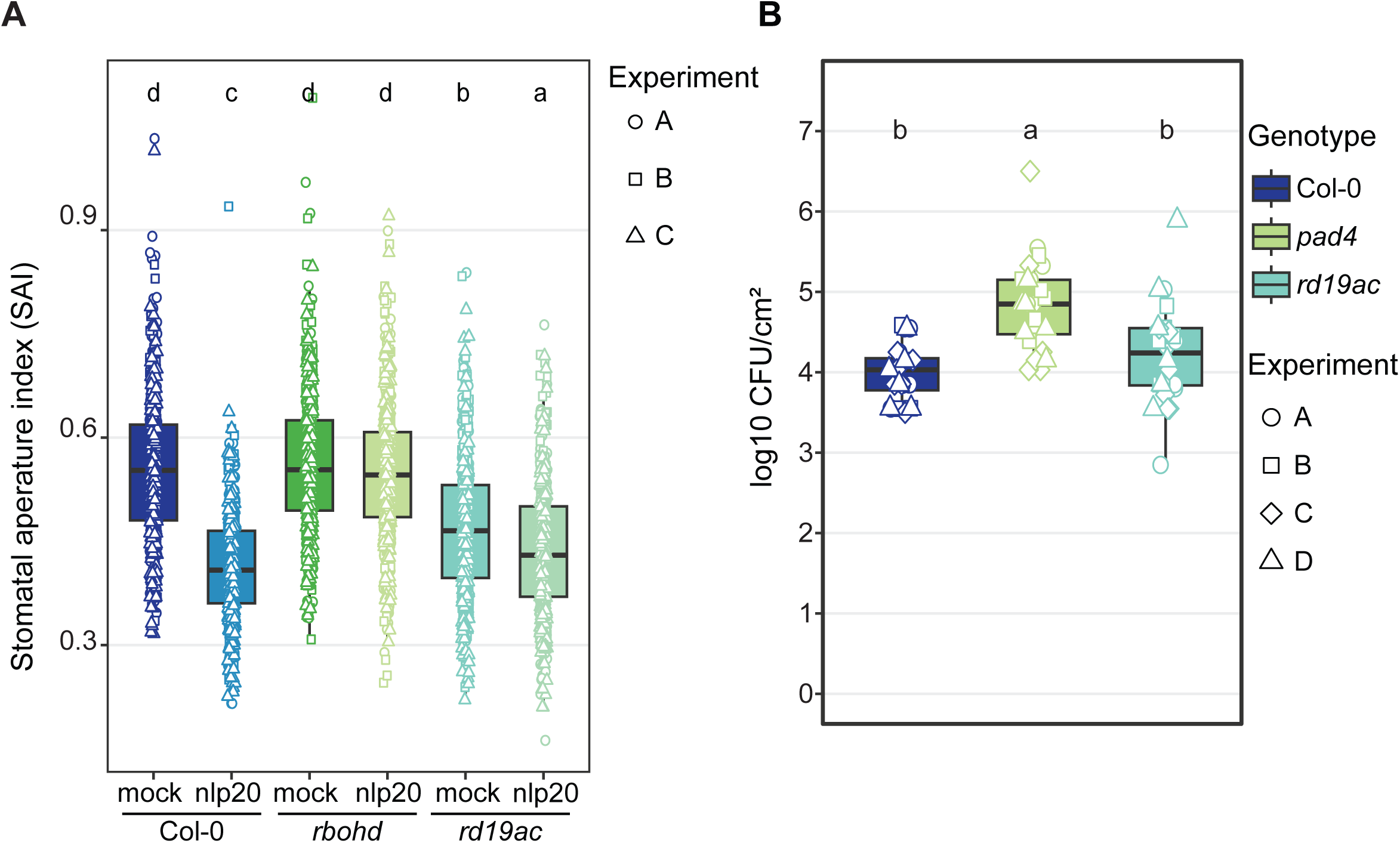
Loss of RD19A and C leads to increased stomatal closure and reduced nlp20 responsiveness without affecting pathogen colonization. **A)** Boxplots of stomata aperture index (SAI) generated by calculating stomata width/length following Rhodamine R staining of epidermal peels from 4.5-week-old Col-0, *rbohd* and *rd19ac*, treated with mock and nlp20 solution for 2 h. **B)** Infection of Col-0, *pad4* and *rd19ac* with *Pst* ΔCor. Boxplots show colony forming units (CFU)/cm^2^ at 3 d after infection, with dots of the same shape representing one biological replicate. Genotypes sharing letters above boxplots do not show statistically significant differences (ANOVA and Tukey HSD test, p < 0.05, n=24). Centre lines of boxplots indicate the median, outer edges the 25^th^ and 75^th^ percentiles and whiskers 1.5 x IQR.

To test whether increased *rd19ac* stomatal closure under mock conditions would affect resistance, plants were spray-inoculated with a virulent *Pseudomonas syringae pv. tomato* DC3000 (*Pst* DC3000) bacterial strain lacking the phytohormone coronatine (*Pst* ΔCor), which it produces to suppress plant stomatal closure (Geng *et al*., 2012; Zheng *et al*., 2012). While *pad4* was more susceptible to *Pst* ΔCor, the *rd19ac* mutant behaved like Col-0 (Figure 3B). Therefore, although *rd19ac* exhibited enhanced basal stomatal closure this did not lead to increased resistance to *Pst* ΔCor (Figure 3B). Taken together, our results suggest a negative effect of RD19 on stomatal closure in nlp20-triggered PTI without impacting stomatal-induced resistance to *Pst* ΔCor.

### RD19 is required for nlp20 induced resistance to *Pst* DC3000 and *Botrytis cinerea* infections

The impact of RD19A and RD19C on nlp20-triggered ROS production and RBOHD protein accumulation (Figure 1D, Figure 2A) prompted us to test whether RD19 influences late infection responses. Comparably to spray-infection with *Pst* ΔCor, the *rd19ac* mutant did not display altered basal resistance to virulent *Pst* DC3000 bacteria (Zeng *et al.,* 2024; Supplementary Figure S4A) or the necrotrophic fungus *Botrytis cinerea* (Supplementary Figure S4B). We next tested the role of RD19 in PTI-induced pathogen resistance. A pre-treatment with flg22 or nlp20 renders Col-0 plants more resistant to subsequent pathogen infection as an outcome of PTI (Zipfel *et al*., 2004; Böhm *et al*., 2014). As expected, pre-treatment with either flg22 or nlp20 in Col-0 led to increased resistance to *Pst* DC3000 and *Botrytis cinerea* compared to mock pre-treated plants, while *fls2* and *rlp23* were PTI defective (Figure 4A and Figure 4B). Pre-treatment with flg22 led to similar restriction of *Pst* DC3000 growth in *rd19ac* and Col-0 (Figure 4A). By contrast, nlp20 pretreatment failed to induce resistance to *Pst* DC3000 in *rd19ac*, resembling *rlp23* or *pad4* defects (Figure 4A; Pruitt *et al.,* 2021). A Flg22 pre-treatment enhanced resistance to *B. cinerea* in Col-0 but not in *rd19ac* (Figure 4B). Notably, resistance to *B. cinerea* in Col-0 was enhanced after nlp20 pre-treatment but reduced in *rd19ac* (Figure 4B). Taken together, these results suggest that RD19A and RD19C are necessary for mounting effective nlp20-induced resistance responses to *Pst* DC3000 and *B. cinerea*.

**Figure 4.**
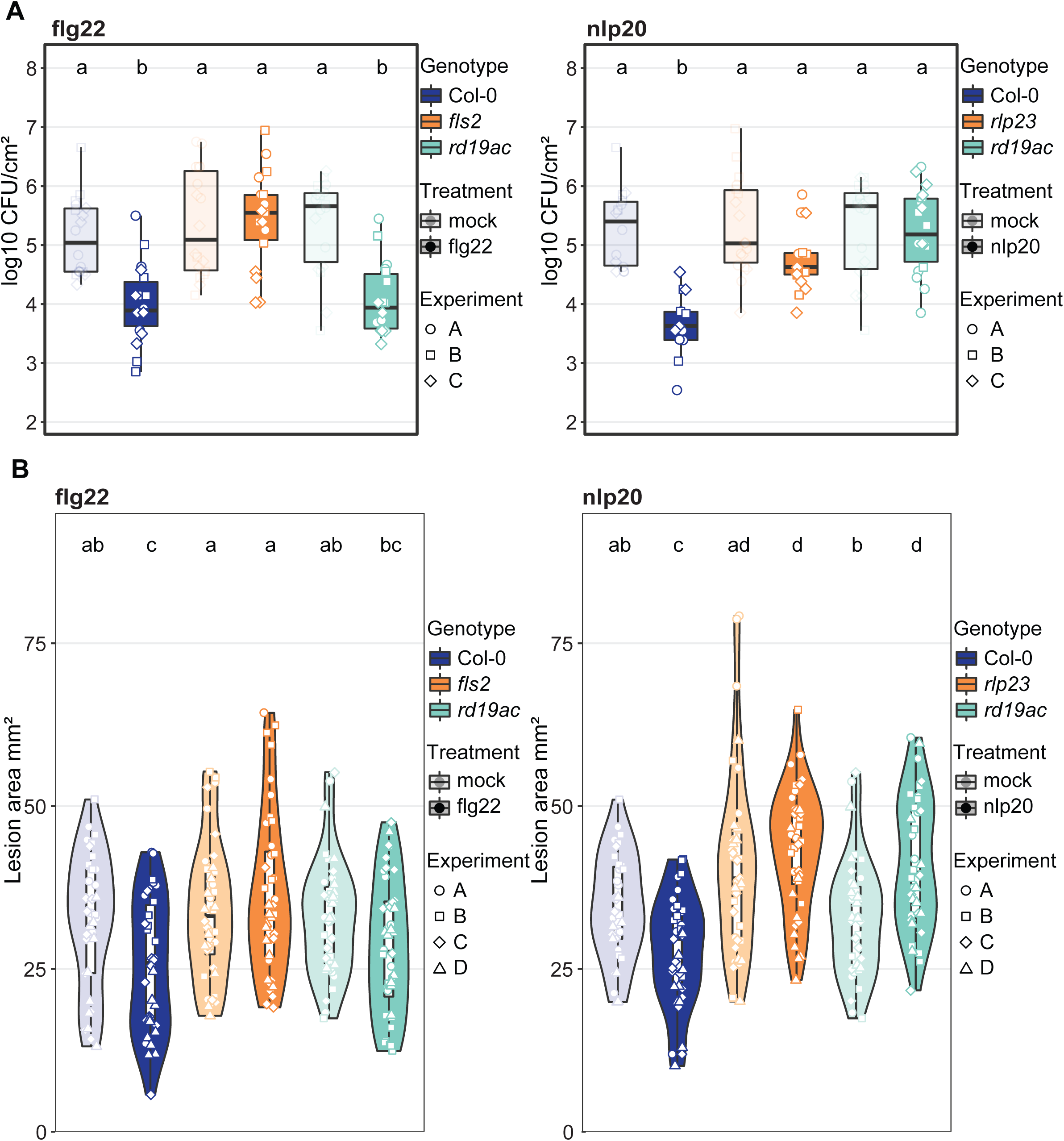
RD19 is required for nlp20 induced immunity to *Pst* DC3000 and *Botrytis cinerea*. flg22 and nlp20 were infiltrated **A)** or sprayed onto leaves **B)** of 4.5-week-old plants and 1 d prior to syringe infiltration with *Pst* DC3000 **A)** or drop inoculation with *B. cinerea* **B)**. Pre-treatment with a mock solution was used as control and is plotted in transparent colours. **A)** Boxplots show colony forming units (CFU)/cm^2^ at 3 d after infection, with dots of the same shape representing one biological replicate. Genotypes sharing letters above boxplots do not show statistically significant differences (ANOVA and Tukey HSD test, p < 0.05, n=18). **B)** Violin blots show lesion areas in mm^2^ with dots of the same shape representing one biological replicate. Genotypes sharing letters above boxplots do not show statistically significant differences (Kruskal-Wallis and pairwise Wilcox-test, p < 0.05, n=35). Centre lines of boxplots indicate the median, outer edges the 25^th^ and 75^th^ percentiles and whiskers 1.5 x IQR.

### Identification of RD19 candidate substrates using High-efficiency Undecanal-based N-Termini EnRichment

Having established that RD19 impacts RBOHD generated ROS upon nlp20 elicitation and contributes to nlp20-induced resistance to a hemibiotrophic and a necrotrophic pathogen, we next aimed to identify RD19A substrates at early (10 min) and late (3 h) time points after nlp20 treatment. RD19A was previously shown to be an active cysteine protease enzyme which might, like other cysteine proteases, cause quite subtle changes to the abundance, size or sequence of other proteins (Demir *et al*., 2018). We therefore employed proteomics-based N-terminomic profiling by High-efficiency Undecanal-based N-Termini EnRichment (HUNTER) (Weng *et al*., 2019). Here, free primary amines, including those at processed protein N-termini generated by endopeptidases like RD19, are modified by reductive methylation and then identified by mass spectrometry, while internal peptides arising from subsequent tryptic proteome digest are depleted (Figure 5A). We compared total extracts of Col-0 and *rd19ac* plants treated with nlp20 for 10 min or 3 h and included untreated samples of both lines as a baseline reference. Across four biological replicates per condition, we identified 7116 unique N-terminal peptides that were quantified in at least three of the four replicates of each condition or uniquely found in one genotype in one condition (Supplementary Table S2). To identify RD19-generated substrates, we searched for N-terminal peptides present only or with significantly higher abundance in Col-0 compared to *rd19ac* (limma-moderated t-test p-value <0.05 with at least 50% increase) that were di- or monomethylated, indicating free proteolysis-generated N-termini *in planta*, and mapped to non-canonical positions within the protein sequence. Applying these criteria identified 106 N-terminal peptides from untreated samples (Figure 5B), 84 peptides at 10 min after nlp20-treatment (Figure 5C) and 76 N-terminal peptides after 3 h of nlp20 treatment (Figure 5D). A Venn diagram of N-terminal peptides identified in Col-0 at the three timepoints revealed very little overlap across the three conditions (Figure 5E). One peptide was shared between untreated and 10 min nlp20-treated samples, representing a likely indirect cleavage of the photosystem II recreation centre protein C (PsbC) before position Tyr11 (Figure 5E). One peptide originating from RD19A with a cleavage position at Asp123 was shared between the 10 min and 3 h nlp20 treatments, while a peptide generated from RD19A cleavage before Leu120 was present across all conditions (Figure 5E; Supplementary Table S2). This latter peptide indicated a cleavage near the RD19A pro-peptide cleavage site, suggesting that RD19A is present in an active form in Col-0 wild type across all three conditions (Figure 5E). Together, these results enabled identification of RD19-dependent candidate substrates cleaved at early and late nlp20-induced immune response stages. The results further show that while RD19 remained active across all three conditions, there was very little overlap of RD19 substrates, highlighting potentially dynamic RD19 substrate targeting during nlp20-mediated immune activation.

**Figure 5.**
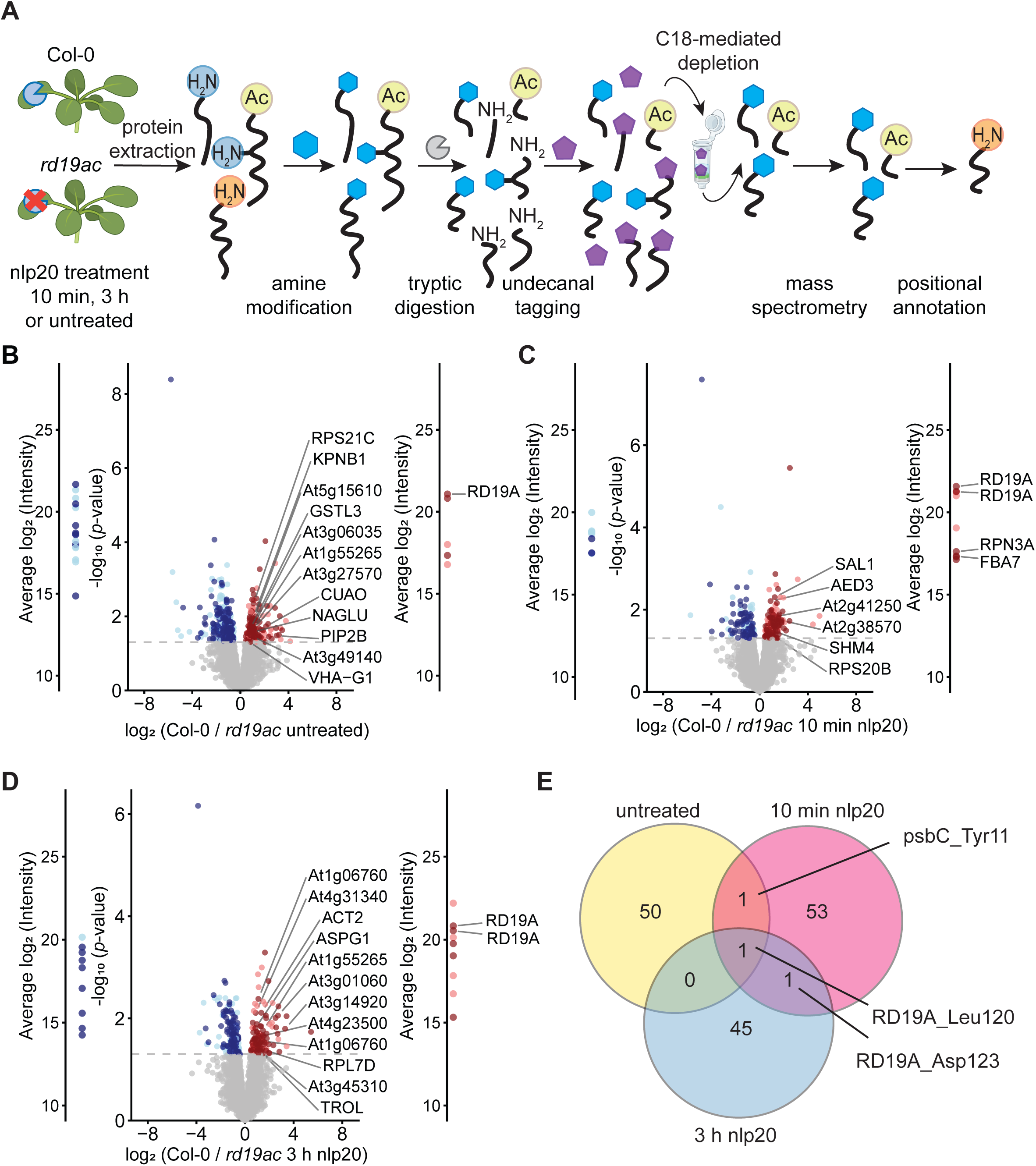
N-terminomics identifies RD19 candidate substrates with roles in immunity. **A)** Schematic overview on sample processing for HUNTER. BioRender.com was used to generate graphics of *Arabidopsis* plants and the column tube. **B) to D)** Volcano plots of N-terminal peptides identified in **B)** untreated plants, **C)** 10 min after nlp20 treatment and **D)** 3 h after nlp20 treatment. N-terminal peptides with significant accumulation (Limma-moderated Student’s t-test p-value < 0.05 and 50% change in abundance) in Col-0 are coloured red, those accumulating in *rd19ac* are blue. N-terminal peptides exclusively found in one genotype are marked on log_2_ intensity scales on either side. **E)** Venn diagram of mono- and di-methylated peptides across the three conditions.

### Proteomic Identification of Cleavage Sites reveals RD19A cleavage preference for non-polar amino acids

To further narrow down HUNTER identified candidates, we aimed to filter the dataset for RD19A direct substrate cleavage signatures. As little experimental information on the amino acid cleavage preferences of RD19 proteases was available, we performed Proteomic Identification of Protease Cleavage Sites (PICS) to determine the sequence specificity of RD19A (Schilling *et al*., 2011; Demir *et al*., 2022*a*). We focused on RD19A because we found consistent activity across all conditions in HUNTER (Figure 5E) and RD19A expression was the most upregulated upon nlp20 treatment compared to the other isoforms (Supplementary figure S1A). RD19A wild-type (*35S*::*RD19A^WT^*-3xHA) or an RD19A catalytically inactive mutant (*35S*::*RD19A^C156A_C159A^*-3xHA) were transiently expressed in *Nicotiana benthamiana* and apoplastic fluid (AF) was isolated. Immunoblotting revealed bands at three different sizes for *35S*::*RD19A^WT^*-3xHA expression, likely corresponding to full length RD19A (45 kDa), proRD19A (42 kDa) and mature RD19A (29 kDa), while expression of *35S*::*RD19A^C156A_C159A^*-3xHA led to one band matching the size of full length RD19A (Figure 6A). AFs containing mutant or wild-type RD19A, or wild-type RD19A pre-incubated with the PLCP inhibitor E64, were then incubated with either trypsin- or GluC-generated *E. coli* proteome-derived peptide libraries for 1 h (Figure 6B). Reactions were stopped by the addition of E64 to all samples and peptide N-termini modified by dimethylation were identified by quantitative data-independent acquisition (DIA) mass spectrometry. We identified 234 unique cleavage sites in the trypsin-library and 70 in the GluC-library detected after incubation with active RD19A compared to catalytically inactive RD19A^C156A/C159A^ and active RD19A pre-incubated with E64. Alignment of the RD19A-generated cleavage sites revealed a strong enrichment of small, aliphatic amino acids such as Ile, Val, Ala and Thr at the P2 position in both datasets (Figure 6C and 6D). Large hydrophobic amino acids (Leu, Tyr, Phe) and polar amino acids (Met, Gln and His) were preferred at P1 and an additional preference for Val and Ala at P4 was observed (Figure 6C and 6D). These results are in line with previously reported cleavage preferences of PLCPs in plants, with nonpolar amino acids showing high prominence in position P2 of the substrate (Petushkova *et al*., 2026).

**Figure 6.**
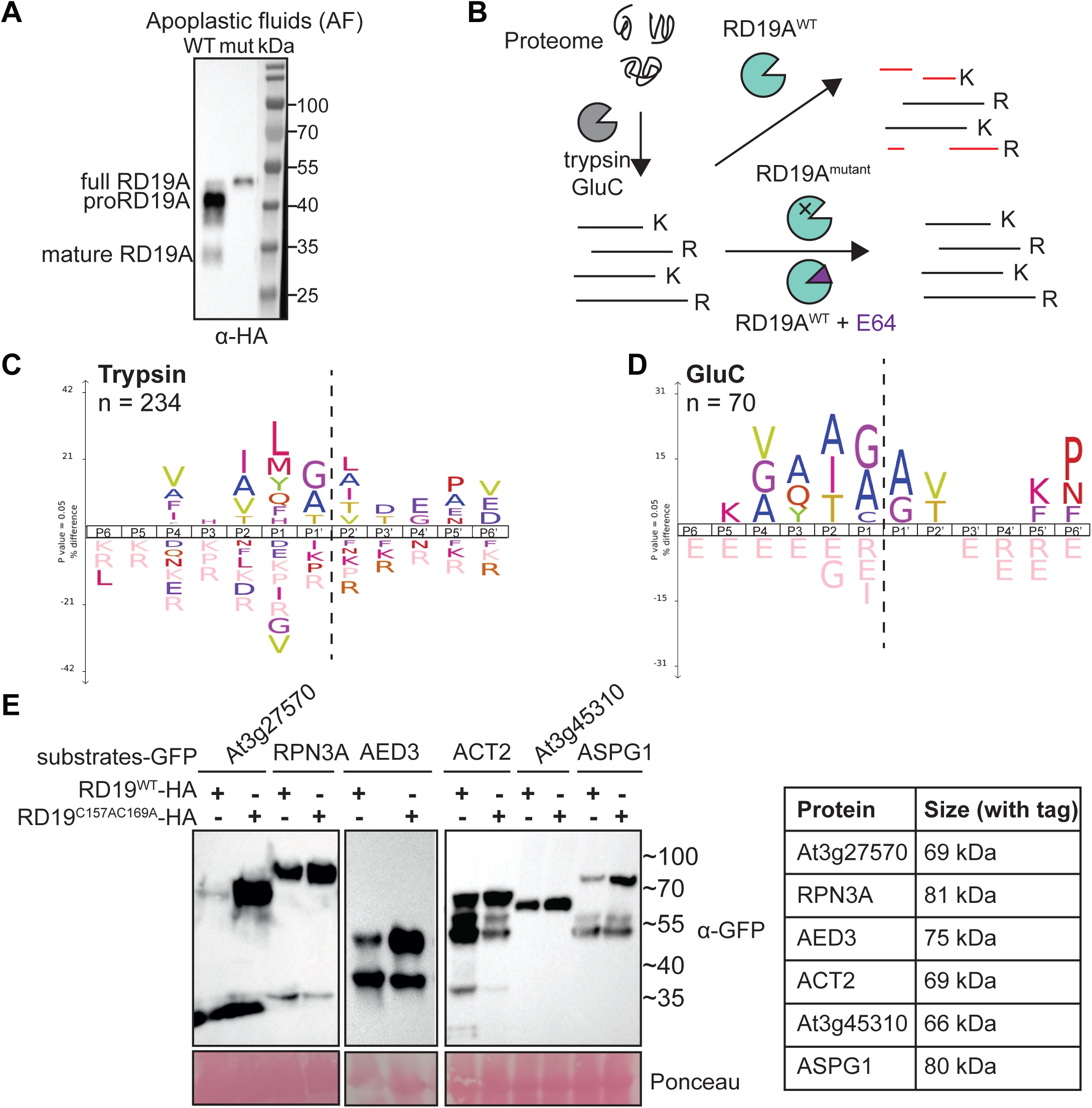
PICS assay reveals RD19A cleavage preferences for non-polar amino acids and prioritizes targets for transient expression analysis. **A)** Immunoblot of apoplastic fluids (AF) from 35S::RD19A^WT^-HA (=WT) and 35S::RD19A^C156A_C159A^-HA (=mut; catalytic mutant), probed with α-HA antibodies. Different maturation forms of RD19A are indicated next to the blot. **B)** Schematic overview on PICS assay workflow. *E. coli* proteins were digested with trypsin or GluC (not shown) and the resulting peptide libraries incubated with RD19A^WT^ (35S::RD19A^WT^-HA) and genetically (35S::RD19A^C156A_C159A^-HA) or chemically inactivated (RD19A^WT^ + E64) controls. Cleavage products of peptides cleaved by RD19A^WT^ but not the controls after 1 h of incubation were identified by quantitative mass spectrometry. **C)** and **D)** iceLogos summarizing **C)** 234 cleavage sites identified with the trypsin peptide library and **D)** 70 cleavage sites identified with the GluC peptide library. The cleaved peptide bond is indicated by the dashed line; the y-axis shows significantly over- or under-represented amino acids compared at each site (p-value < 0.05). **E)** Immunoblotting of substrate candidates after co-expression with 35S::RD19A^WT^-HA or 35S::RD19A^C156A_C159A^-HA in *N. benthamiana.* Substrate candidates were YFP tagged and detected using α-GFP antibodies. Ponceau-stained Rubisco served as a loading control, to ensure equal protein levels across samples. Images of different blots shown are representative of protein levels and cleavage patterns observed in three independent biological replicates.

### *In planta* validation of RD19A substrates with previously described roles in immunity

We next integrated the cleavage preferences revealed by PICS with the list of candidate cleavage sites identified in our HUNTER N-terminome analysis to identify direct putative RD19A substrates (Table 1 and Figure 5B-D). Peptides derived from proteins with annotated localization in chloroplasts or mitochondria were excluded, as RD19A has predicted or reported subcellular localization in the apoplast, endosomes/vacuole, cytosol and nucleus (Bernoux *et al*., 2008; Richau *et al*., 2012; Zeng *et al*., 2024). This analysis produced 35 N-terminal peptides, of which six were derived from cleavages with a Val, Ile, Ala or Thr at P2, consistent with the PICS cleavage motif (Table 1; Figure 6C and 6D). We focused specifically on a match with the P2 cleavage site because this position was reported to be important for cysteine protease specified cleavage (Petushkova *et al*., 2026). A literature survey revealed additional potential RD19 substrates with previously described roles in immunity among HUNTER candidates (Table 1). For validation *in planta*, we tested a total of six HUNTER identified substrates, four of which showed a match with the predicted PICS motif (At3g27570, 26S proteasome non-ATPase regulatory subunit 3 homolog A (RPN3A), ACTIN2 (ACT2) and Thiol protease aleurain-like At3g45310) and two which were part of the identified proteins with described roles in immunity (APOPLASTIC EDS1-DEPENDENT 3 (AED3) and ASPARTIC PROTEASE IN GUARD CELL 1 (ASPG1/AED2)) (McDowell *et al*., 1996; Beers *et al*., 2004; Yang *et al*., 2004; Loreti *et al*., 2005; Yao *et al*., 2012; Breitenbach *et al*., 2014).

**Table 1:**
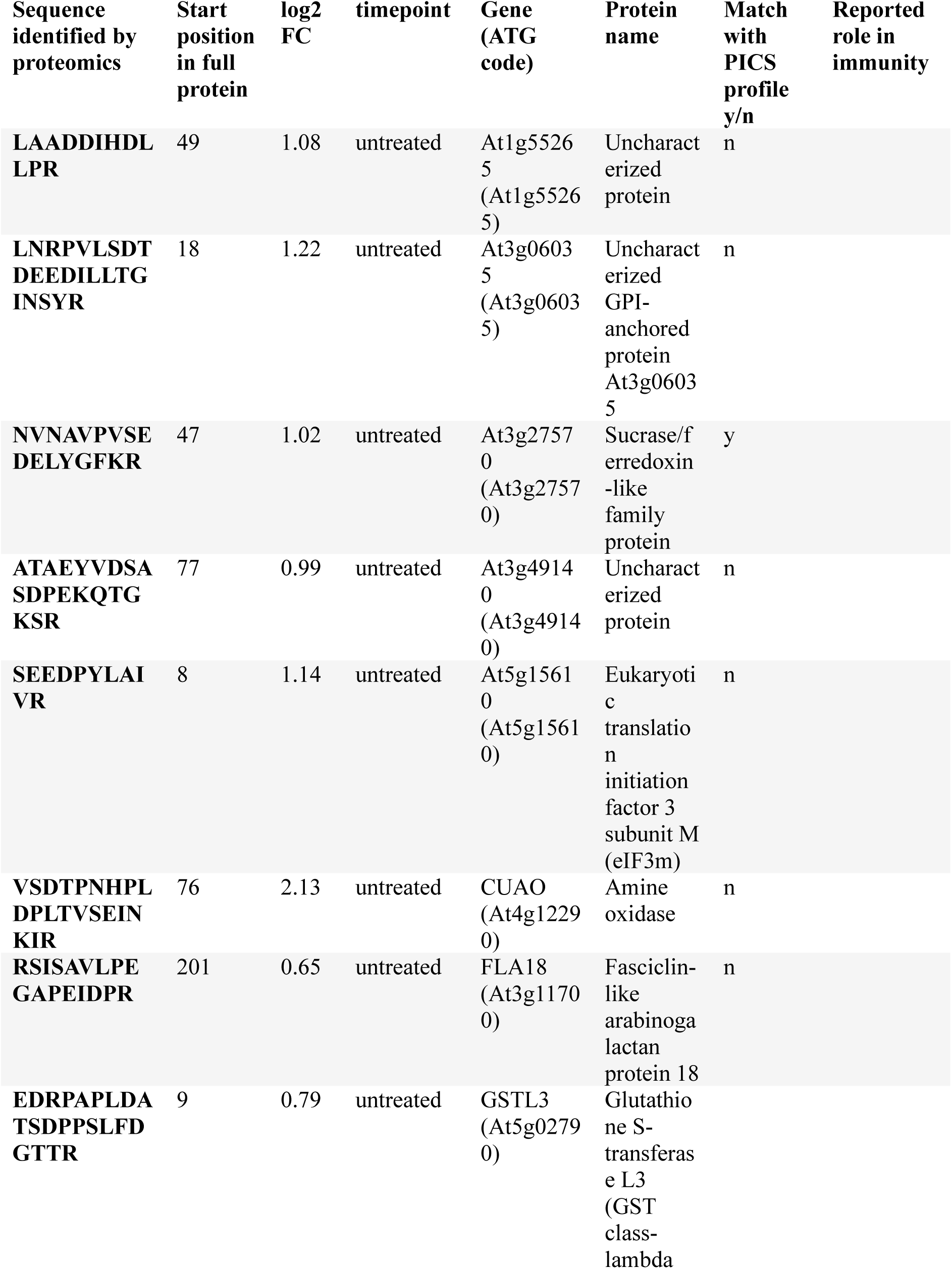
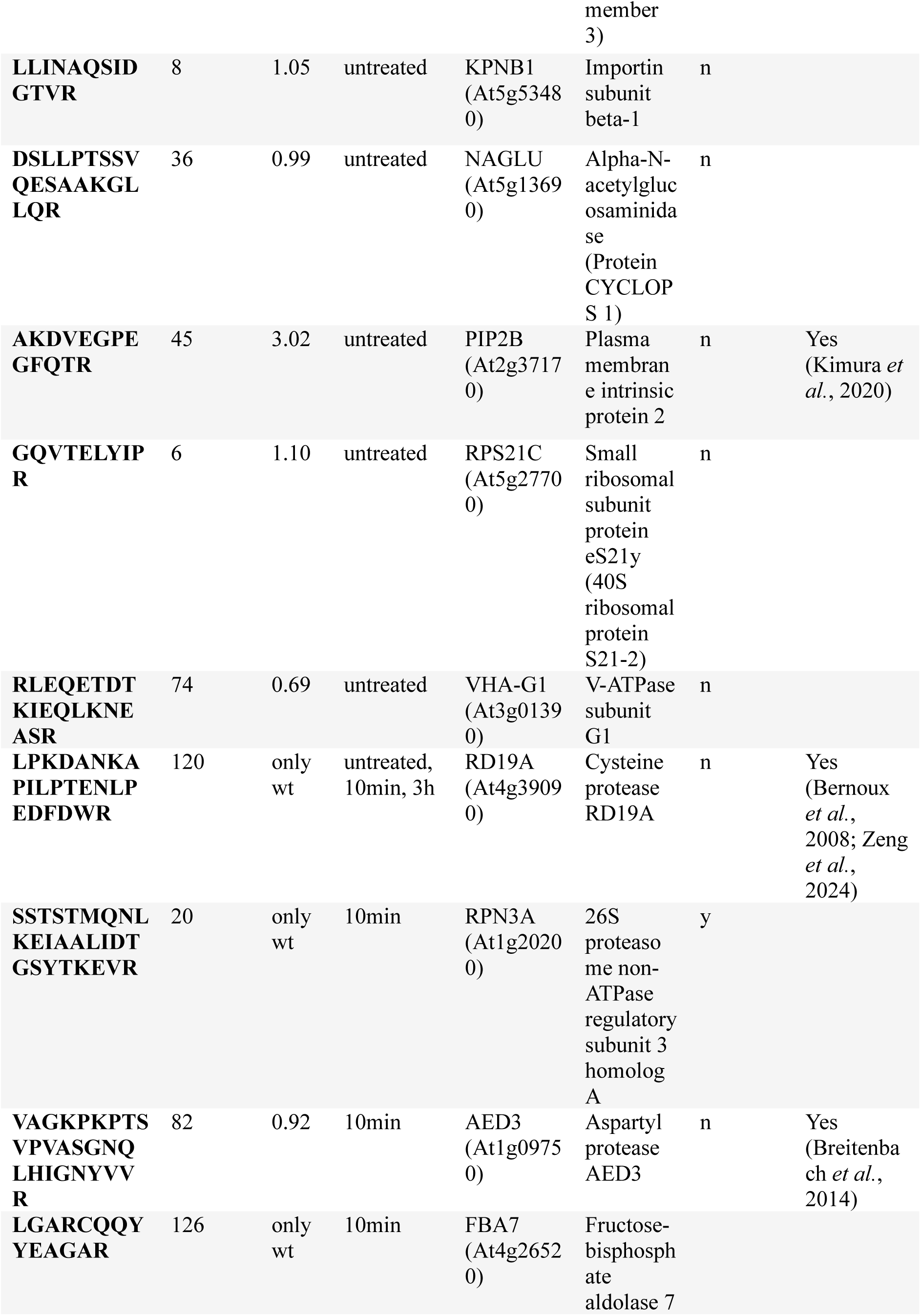
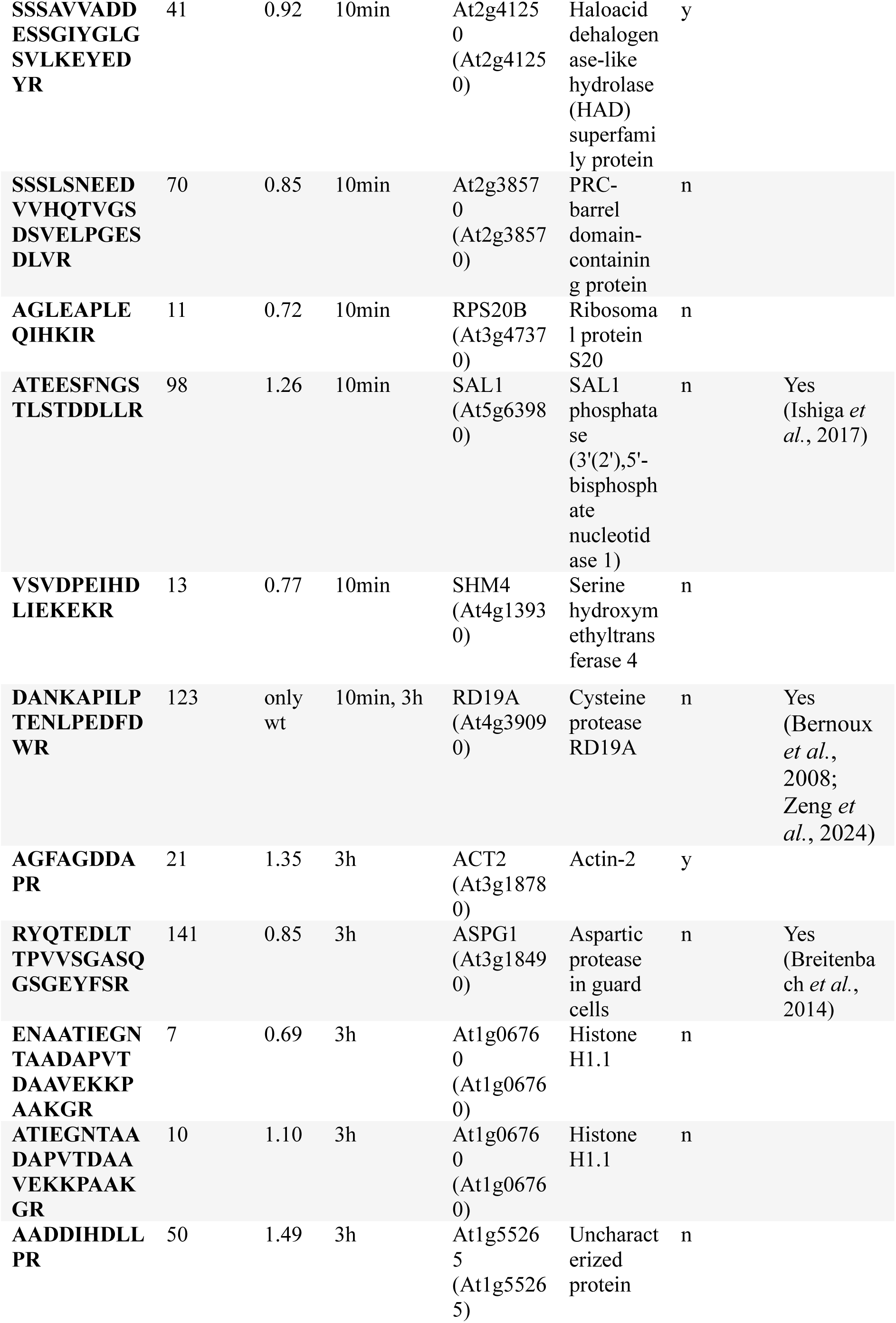
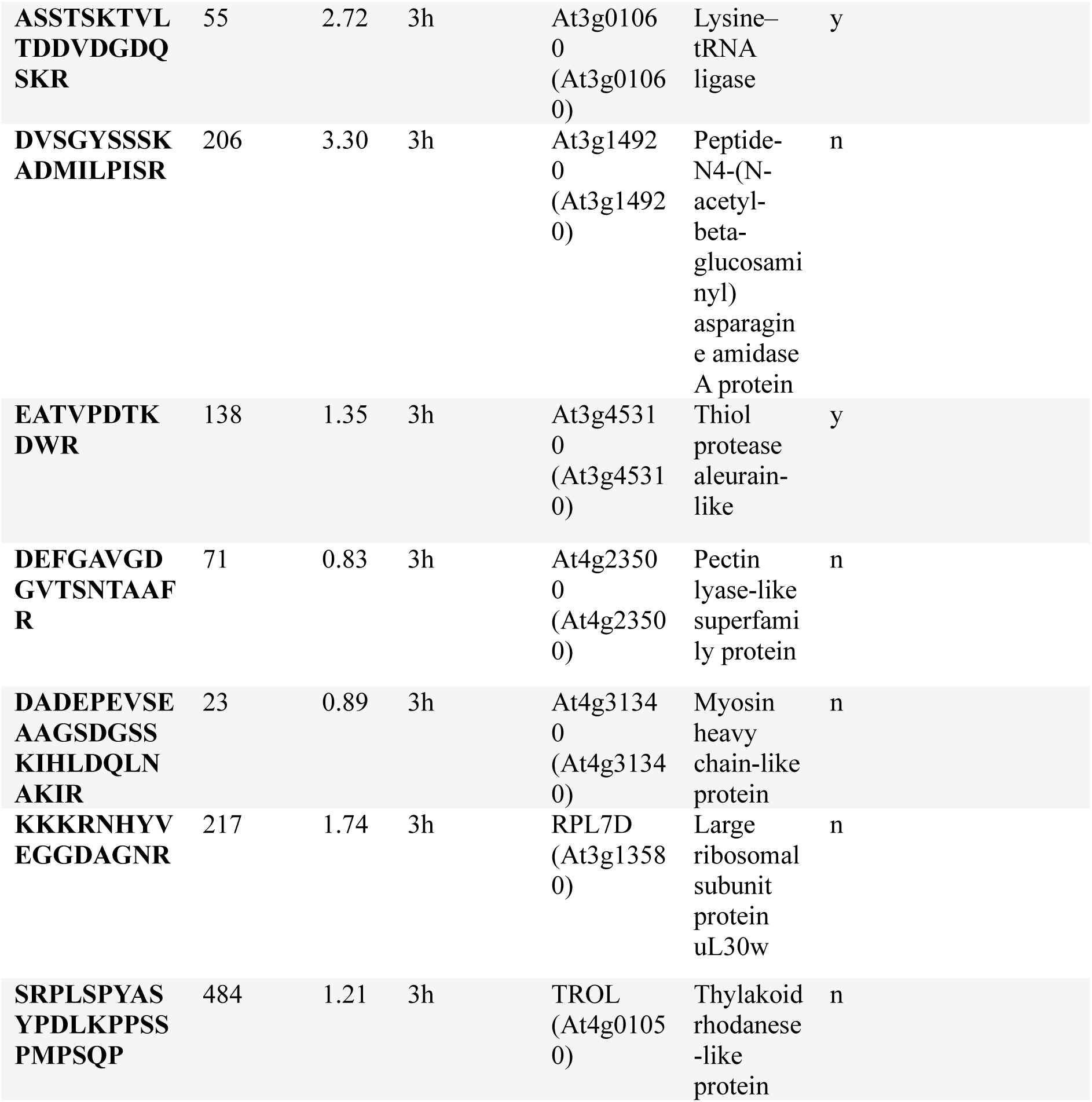
RD19 candidate substrates identified by HUNTER. Match with PICS data is indicated by yes/no (y/n). References to reported roles in immunity are listed in the last column.

To verify substrate cleavage by active RD19A *in planta* we transiently overexpressed the six candidate proteins tagged with YFP together with active or inactive *35S::RD19A*-3xHA in *N. benthamiana* (Figure 6E). Co-expression with active RD19A compared to inactive RD19A (RD19A^C156A/C159A^) revealed changed band patterns and reduced band intensity of the full-length protein size band in substrate At3g27570 (identified in non-treated conditions), AED3 (identified in HUNTER 10 min after nlp20-treatment) and ACT2 and ASPG1 (detected 3 h after nlp20-treatment) (Figure 6E). This result suggested that lower amounts of the full-length target proteins and increased levels of lower molecular weight bands were present due to RD19A protease activity. At3g27570-YFP (69 kDa) showed one additional band ∼35 kDa; however, this did not correspond to the generated cleaved protein starting at position 47, based on the N-terminomics peptide analysis (Figure 6E and Table 1). By contrast, ACT2, AED3 and ASPG1 all showed additional bands at protein sizes which appeared to correspond to cleavage detected by HUNTER (Supplementary Table S2; Figure 6E). We were especially interested where cleavage occurred for aspartic proteases AED3 and ASPG1, since these could be indicative of an activating or inactivating processing. For AED3, the HUNTER identified peptide spans amino acids 82 to 107 (Table 1; Supplementary Table S2) indicating cleavage before amino acid 82 at the AED3 N-terminus. Notably, AlphaFold3 modelling (Abramson *et al*., 2024) of AED3 predicted AED3 cleavage prior to the protease domain (with two aspartic acid active sites in position D122 and D328, highlighted in green) and increased confidence in the structural prediction of the cleaved version (from pTM = 0.77 to pTM = 0.92) (Supplementary Figure S6A). A ∼66 kDa lower molecular weight band observed on Western blots matched the 39 kDa size of cleaved AED3 with YFP tag (Figure 6E). Superimposing the models of full length and cleaved AED3 supported removal of the pro-domain and preservation of D122 and D328 active sites with an RMSD score of 0.446 Å (Supplementary figure S6B). By comparison, ASPG1 was cleaved prior to amino acid 141 (Table 1; Supplementary Table S2). AlphaFold3 modelling suggested cleavage of the pro-domain prior to the protease domain with asparagine active sites at positions 180 and 379 (Supplementary figure S6C) and the molecular weight of cleaved ASPG1 (36 kDa) matched the tagged form observed on western blots (Figure 6E). Here, superimposing full length and cleaved ASPG1 suggested that the pro-domain is located on top of the active site in full length ASPG1, which might highlight importance of proteolytic cleavage for active site availability (RMSD score of 1.011 Å; Supplementary Figure S6D).

In conclusion, our proteomic data analysis and transient assay validations identify several RD19 substrates in nlp20-triggered immunity. Of these, two modelled aspartic proteases display probable RD19A protease-dependent activating cleavage.

## Discussion

The plant immune system requires tight regulation to prevent over stimulation. Proteolysis plays an important role in this regulation by cleaving substrates for activation or inactivation (van der Hoorn and Jones, 2004; Misas-Villamil *et al*., 2016). The cysteine protease RD19 was previously described as a component of ETI and basal immunity and an interactor of PAD4 but not SAG101 or EDS1 (Bernoux *et al*., 2008; Zeng *et al*., 2024). As the EDS1-PAD4-ADR1 branch is an ETI-PTI signalling convergence point (Dongus and Parker, 2021; Pruitt *et al*., 2021; Fliegmann *et al*., 2026), in this study we investigated whether RD19 plays a role in PTI and, if so, whether that mirrors the contribution of PAD4 in defence potentiation. Here we found that while the role of RD19 does not align closely with that of PAD4, RD19 is required for the regulation of ROS in response to nlp20, likely by controlling RBOHD protein accumulation (Figure 1D and Figure 2A). We further established that RD19 contributes to mounting robust PTI defence and speculate this could be through direct or indirect control of RBOHD protein turnover and a proteolytic cascade involving two different aspartic proteases as shown in a model (Supplementary Figure 7).

RBOHD was previously characterized as a major ROS production and defence hub during PTI and ETI (Kadota *et al*., 2015; Ngou *et al*., 2021, 2022; Yuan *et al*., 2021), and is thus subject to multiple levels of regulation mediated principally by Ca^2+^-dependent and -independent protein kinase phosphorylation (Dubiella *et al*., 2013; Kadota *et al*., 2014; Li *et al*., 2014). We found that an increased ROS burst upon nlp20-treatment in *rd19ac* mutants coincided with increased accumulation of RBOHD protein but not gene expression (Figure 2A and 2B and Supplementary Figure S2). This suggests that the RD19A and RD19C cysteine proteases limit ROS accumulation, at least in part, by controlling RBOHD turnover. Consistent with an increased accumulation of RBOHD in untreated *rd19ac* plants, stomatal closure was higher under mock conditions compared to Col-0 (Figure 3A). However, *rd19ac* stomata displayed decreased stomatal closure responsiveness than Col-0 when exposed to nlp20 (Figure 3A). While RBOHD was the main driver of the observed increased ROS burst in *rd19ac* (Figure 2C and Figure 2D), stomatal ROS is generated by both RBOHD and RBOHF (Torres *et al*., 2002; Kwak *et al*., 2003; Bright *et al*., 2006) and notably, RBOHF is the major regulator of stomatal ROS production in PTI (Arnaud *et al*., 2023). It is therefore possible that general mis-regulation of RBOHD generated ROS in *rd19ac* interferes with RBOHF function and stomatal responsiveness to nlp20 (Figure 3A).

A previous study showed that RBOHD is degraded inside the plant vacuole by the cysteine protease XCP1 (Liu *et al*., 2023). This occurred under resting (non-triggered) conditions, following RBOHD ubiquitination and phosphorylation driven vacuolar targeting (Lee *et al*., 2020; Liu *et al*., 2023). With immune activation, XCP1 was inhibited by CYSTATIN 6 (CYS6), allowing RBOHD recycling to the plasma membrane (Lee *et al*., 2020; Liu *et al*., 2023). A comparable process might apply to RD19 in this study, since RD19 regulated RBOHD accumulation in resting and nlp20-stimulated plants (Figure 2A and 2C). We hypothesized that RBOHD itself could be a target of RD19 (Supplementary Figure 7). However, the N-terminomic profiling did not identify RBOHD, suggesting that the RD19 regulatory effect is indirect (Figure 5B-D/Table 1). In line with this hypothesis, two of the proposed RD19 substrates, *Arabidopsis* ASPG1/AED2 and ACT2 (Figure 6E), were previously described as regulators of ROS production (Yao *et al*., 2012; Takáč *et al*., 2024). Overexpression of ASPG1 led to reduced levels of H_2_O_2_ by increasing ROS scavenging activities of superoxide dismutase and catalase, leading to improved drought tolerance (Yao *et al*., 2012). Mutants of ACT2 (*der1-3*) had increased abundance of PM aquaporins PLASMA MEMBRANE INTRINSIC PROTEIN1 (PIP1) and PIP3 in roots, explaining their increased drought stress resistance compared to wild-type plants (Takáč *et al*., 2024). It remains to be tested whether ASPG1/AED2 or ACT2 are functionally related to RBOHD or if cleavage by RD19A impacts their activities. Previous studies of RD19 described a localization in the vacuole and mobile vacuole-associated compartments (Bernoux *et al*., 2008; Zeng *et al*., 2024). It is possible that RBOHD turnover in the vacuole occurs rapidly and we were therefore unable to detect RBOHD derived vacuolar peptides in our N-terminomic profiling. Indeed, the RD19 pH activity spectrum (Richau *et al*., 2012) indicates that RD19 should be active in the vacuole and apoplast, while our *N. benthamiana* transient expression assays show significant RD19 accumulation and activity in the apoplast (Figure 6A; Supplementary Figure 7). Future experiments are required to test for vacuolar turnover of RBOHD mediated by RD19 or a possible RD19 inhibitor which would control this degradation in a similar manner to XCP1 regulation by CYS6 (Liu *et al*., 2023).

In our PICS analysis, we used RD19A derived from apoplastic fluids after transient overexpression in *N. benthamiana* and identified a cleavage preference for aliphatic amino acids in position P2 prior to the cleavage site (Figure 6C and Figure 6D). These findings are in line with previous reports of PLCP cleavage preferences (Choe *et al*., 2006; Petushkova *et al*., 2026) and further support activity of RD19A in the apoplast. Moreover, numerous putative substrates identified by HUNTER have known or predicted apoplastic localization (Table 1). In our experiments, RD19A was active under all three tested conditions – non-treated samples and at 10 min or 3 h after treatment with nlp20 (Figure 5B-E). Strikingly, substrates identified in the three conditions showed very little overlap, suggesting that RD19A targets different proteins depending on status and timing of the immune response (Figure 6E). We cannot rule out that RD19A changes sub-cellular location and is active in different compartments under changing immune conditions. Indeed, we previously found that RD19 co-localizes to the nucleus with PAD4 (Zeng *et al*., 2024). It is not known whether RD19 is active in the nucleus, as proposed for its nuclear co-localization with PopP2 (Bernoux *et al*., 2008) or if nuclear localization dampens RD19 activity, since it would be outside its proposed pH activity spectrum (Richau *et al*., 2012).

Establishing appropriate spatiotemporal immune responses is essential for plants to secure full resistance during infections (Zhu *et al*., 2023). Full resistance requires PTI and immune activation in systemic tissue, both relying on the EDS1-PAD4-ADR1 node and the production of ROS (Wittek *et al*., 2014; Pruitt *et al*., 2021; Mittler *et al*., 2022; Feehan *et al*., 2023; Cao *et al*., 2024; Fliegmann *et al*., 2026). We found that *rd19ac* mutants were defective in nlp20-induced resistance to *Pst* DC3000 and *Botrytis cinerea* (Figure 4). This could be caused in part by a disbalance of ROS levels which are necessary to establish an effective spatiotemporal immune response and resistance to subsequent pathogen infection (Figure 2A and 2B; (Cao *et al*., 2024). The removal of RD19 might deregulate a PAD4-dependent feed-forward loop generated by RBOHD ROS and amplified cell-to-cell by other immune components such as CPKs (Dubiella *et al*., 2013), thus explaining PAD4-dependency of the increased ROS burst in *rd19ac* plants (Figure 1D).

In SAR certain levels of mobile H_2_O_2_ were found to be required to sulphenylate the transcription factor CCA1 HIKING EXPEDITION (CHE) which, in turn promoted SA-biosynthesis (Cao *et al*., 2024). ROS above a certain level inactivated CHE and broke the signal amplification loop needed to establish SAR (Cao *et al*., 2024). Since ROS is also essential for induced resistance, we speculate there is a similar disruption present in *rd19ac* (Figure 4; Supplementary Figure 7). Notably, RD19A was found to be a positive regulator of SAR and a target of the SAR negative regulator miR159b (Kumar *et al*., 2025). The expression of *RD19A* and other miR159b targets was induced upon SA treatment and required intact SA signalling, suggesting a positive feedback mechanism, also depending on EDS1-PAD4, which promotes SA biosynthesis (Cui *et al*., 2017; Kumar *et al*., 2025).

The identified RD19 aspartic protease substrates ASPG1/AED2 and AED3 (Figure 6) were first detected in a proteomic analysis of apoplast-enriched extracts to screen for proteins involved in EDS1-PAD4 dependent SAR (Breitenbach *et al*., 2014). While AED1 suppressed SAR (Breitenbach *et al*., 2014) it was not established whether other AED family members, such as AED2 and AED3, identified here, have a similar role. Sequence analysis and AlphaFold3 modelling predicted that both ASPG1/AED2 and AED3 are cleaved by RD19 at their N-termini, which removes the protease pro-domains leading to activation without altering the structure around the aspartyl active sites (Supplementary Figure S6). One possible scenario is that in wild-type, RD19 activates a protease cascade involving cleavage of AED3 and ASPG1 as well as other substrates (at different timepoints), which then become active and modulate nlp20-triggered immunity and defence potentiation (Supplementary Figure 7).

Our work has uncovered a previously unrecognized role for the cysteine protease RD19 as a regulator of RBOHD generated ROS and PTI. This finding coupled with a proteomic-based analysis of RD19A immune-related substrates lays the foundation for future unravelling protease fine-tuning of plant innate immunity.

## Supplementary data

Table S1: Primers used in this study.

Table S2: Proteomics data of RD19 substrate candidates identified as more abundant in Col-0 than *rd19ac*.

Supplementary methods: detailed mass spectrometry and data analysis

Supplementary Figure S1: RD19 expression profile upon PTI activation and its contribution in PTI-induced ROS burst.

Supplementary Figure S2: *RBOHD* gene expression at later timepoints (15 min and 30 min).

Supplementary Figure S3: Cell type specific gene expression of *RD19A* and *RD19C* from publicly available RNAseq dataset (Tang *et al.,* 2023).

Supplementary Figure S4: Role of RD19 in basal immune responses to *Pst* DC3000 and *Botrytis cinerea*.

Supplementary Figure S5: RD19 transient expression in *N. benthamiana*.

Supplementary Figure S6: AlphaFold3 modelling of AED3 and ASPG1.

Supplementary Figure S7: Working model of RD19 activity during nlp20-triggered PTI.

## Acknowledgements

We thank Johana Misas-Villamil at the University of Cologne for sharing the L1 MoClo vector for heterologous expression in *N. benthamiana* and for helpful discussion and support throughout the project. We thank Daniel Moser for sharing the protocol for PICS sample preparation. We also thank past and present members of the Parker lab for support.

## Author contributions

G.H. and J.E.P. designed the research; G.H., M.M., F.L. and J.B. performed experiments; G.H., M.M. and P.F.H. analysed the data; G.H. and P.F.H. prepared figures and final datasets; G.H., F.L., P.F.H. and J.E.P. wrote the manuscript with contributions from all authors.

## Data availability

The mass spectrometry proteomics data have been deposited to the PRIDE repository (Perez-Riverol *et al*., 2025) with the dataset PXD084587 (reviewer access token: mLJa8qobEBo5) for the PICS data and PXD084625 (reviewer access token: GTHbWx8QIxB3) for the HUNTER data. All other data generated or analysed in this study can be found within the manuscript and its supporting materials.

