## Supplementary material for "*Arabidopsis* cysteine protease RD19 limits RBOHD-generated ROS production in pattern triggered immunity": Suppl Figures S1-S7_Suppl Table S1_Suppl methods

### Supplementary data file:

#### Supplementary figures:

#### Supplementary Figure 1

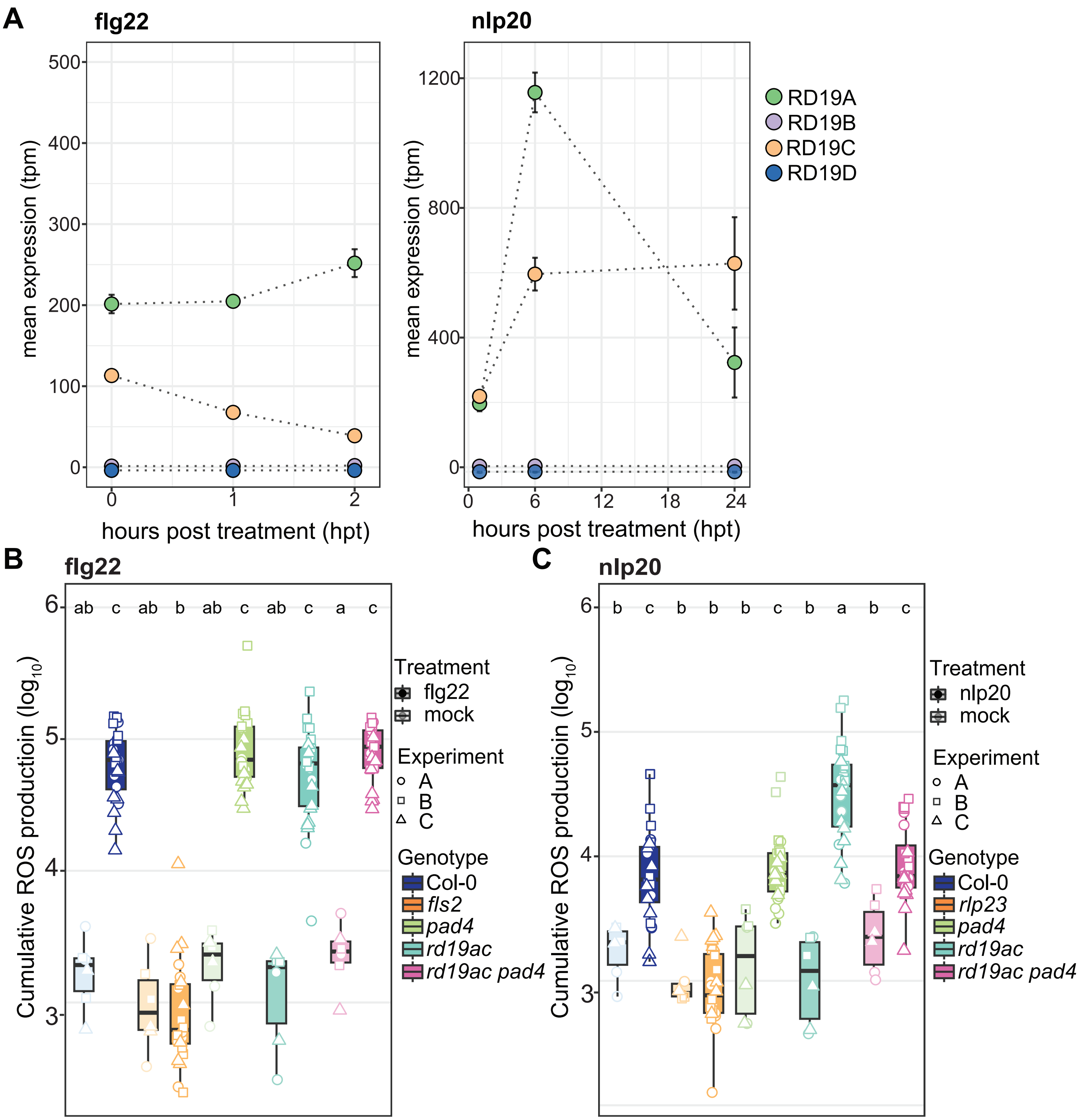


**Supplementary Figure S1. Expression profile of *RD19* genes upon PTI activation and RD19A and RD19C contributions to a PAMP-induced ROS burst**. **A)** *RD19* expression analysis of publicly available RNA-seq data sets following flg22 and nlp20 treatment. RNA-seq data sets were downloaded from the NCBI SRA data bank (flg22: PRJNA339762; GEO: GSE85923, nlp20: PRJNA549850; GEO: GSE133053). Expression of *RD19A*, *RD19B*, *RD19C* and *RD19D* are shown at h post infection (hpi) with flg22 or nlp20. Treatments were performed on 12- and 10-day old Col-0 seedlings, respectively. **B)** and **C)** Area under curve (AUC) for ROS bursts following elicitor treatments with **B)** flg22 and **C)** nlp20. Boxplots show $log_{10}$ transformed AUC values over 60 min ROS measurements with dots of the same shape representing one biological replicate. Genotypes sharing letters above boxplots do not show statistically significant differences (ANOVA and Tukey HSD test, $p < 0.05$, n=24 (flg22 and nlp20) and n=6 (mock) leaf discs from three independent biological replicates). Centre lines of boxplots indicate the median, outer edges the 25th and 75th percentiles and whiskers 1.5 x IQR.

### Supplementary Figure 2

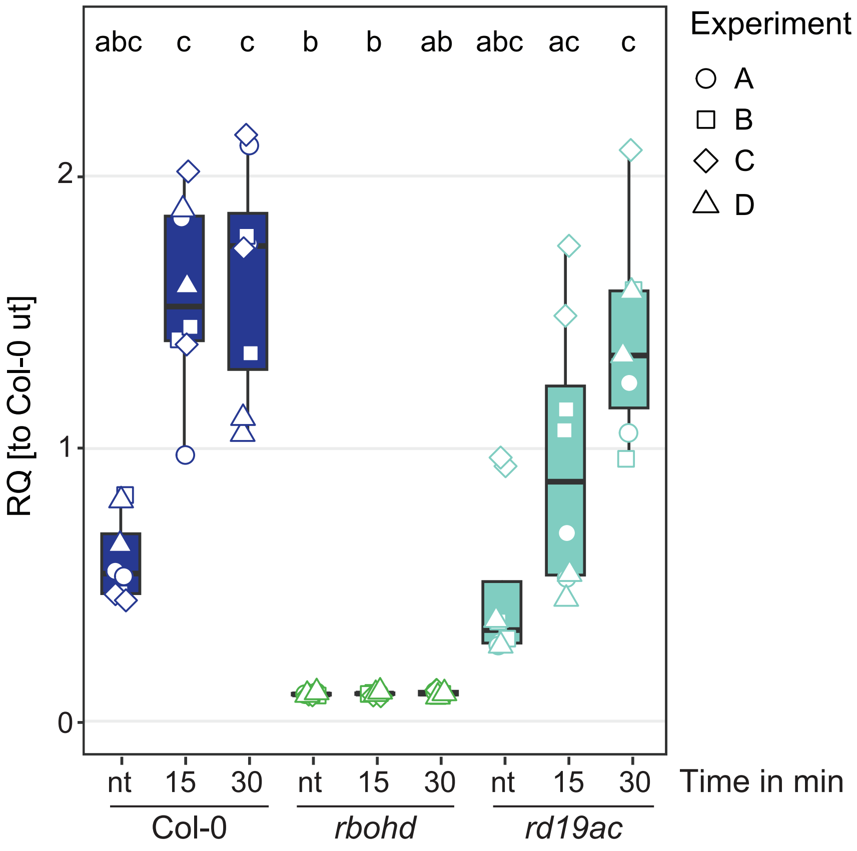


**Supplementary figure S2. Increased RBOHD protein accumulation is independent of *RBOHD* gene expression.** *RBOHD* gene expression changes were measured in non-treated (nt) leaf tissue, and at 15 min and 30 min after nlp20 infiltration. Relative quantification is represented in boxplots with dots of different shapes indicating biological replicates. Data was normalized to Col-0 mock conditions in each biological replicate. *UBI5* was used as a reference gene. Genotypes sharing letters above boxplots do not show statistically significant differences (Kruskal-Wallis and Nemenyi-Test, $p < 0.05$, n=8). Centre lines of boxplots indicate the median, outer edges the 25$^{th}$ and 75$^{th}$ percentiles and whiskers 1.5 x IQR.

### Supplementary Figure 3

Gene expression following infection with *Colletotrichum higginsianum*

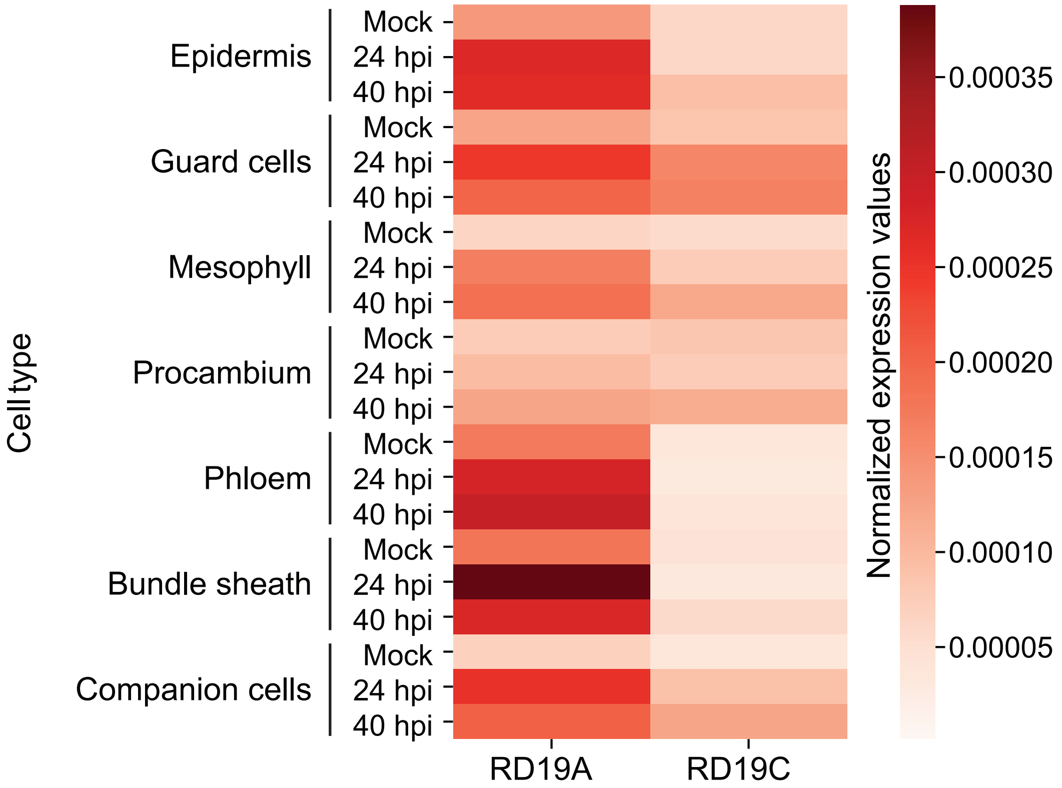


**Supplementary figure S3. Analysis of cell-type specific expression of *RD19A* and *RD19C* from a publicly available single cell RNAseq dataset (Tang *et al.,* 2023) following Arabidopsis Col-0 infection with *Colletotrichum higginsianum* at 24 h and 40 h.** Mock treatment was used as a control. Heatmap depicts normalized expression values and the expression matrix was generated from monocle pipeline by aggregating expression of each gene in cells from the same cell type and normalizing this to cell number as a size factor.

### Supplementary Figure 4

**A**

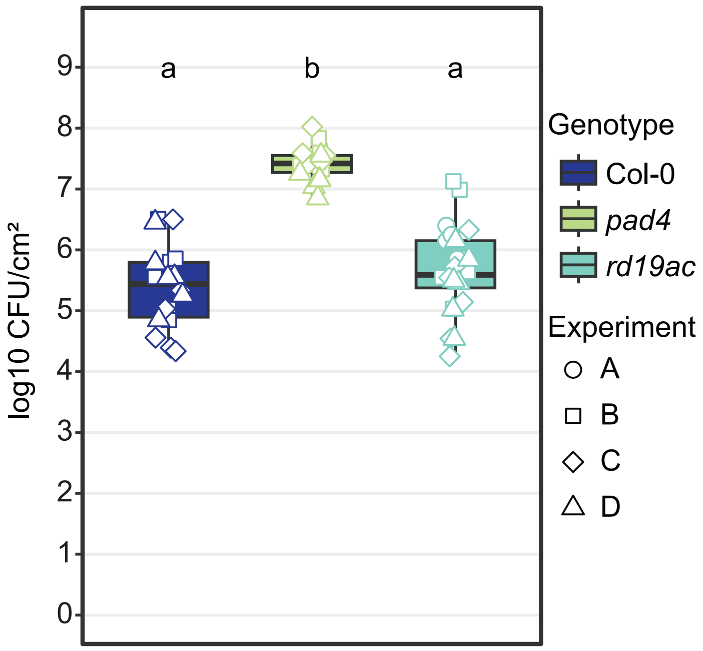


**B**

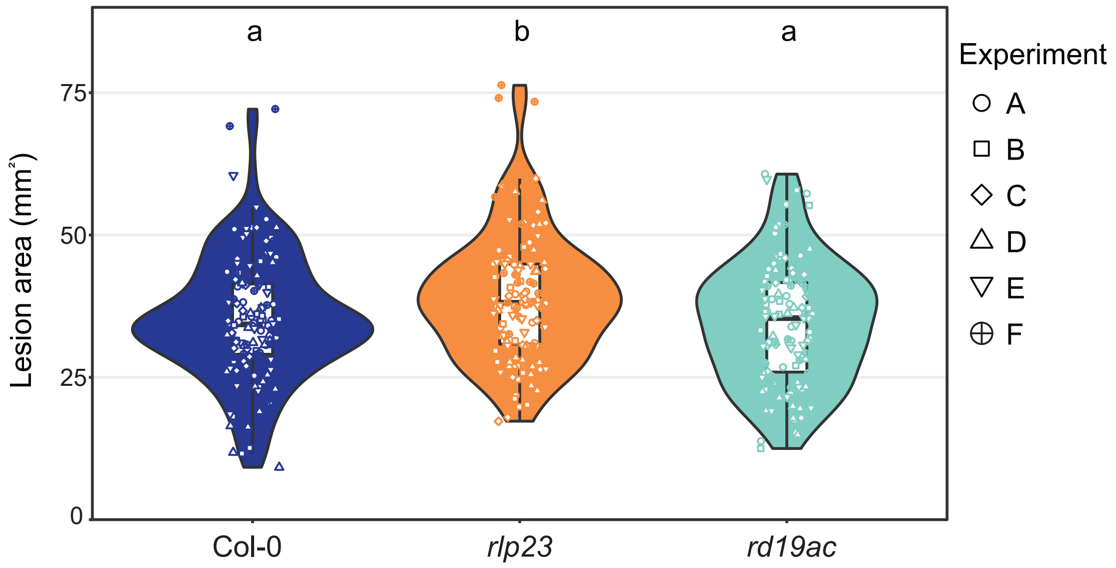


**Supplementary figure S4. RD19A and C are not required for basal immune responses to *Pst* DC3000 and *Botrytis cinerea*. A)** *Pst* DC3000 growth assay in leaves of 4.5-week-old *Arabidopsis* Col-0, *pad4* and *rd19ac* at 3 d post syringe infiltration. Boxplots show colony forming units (CFU)/$cm^2$ with each shape representing one biological replicate. Genotypes sharing letters above boxplots do not show statistically significant differences (ANOVA and Tukey HSD test, $p < 0.05$, n=24). **B)** *B. cinerea* spores were drop-inoculated on leaves of 4.5-week-old *Arabidopsis* Col-0, *rlp23* and *rd19ac* and lesion areas were measured at 2 dpi using ImageJ as a proxy of pathogen growth. Violin blots show lesion areas in $mm^2$, with dots of the same shape representing one biological replicate. Genotypes sharing letters above boxplots do not show statistically significant differences (Kruskal-Wallis and pairwise Wilcox-test, $p < 0.05$, n=90). Centre lines of boxplots indicate the median, outer edges the 25$^{th}$ and 75$^{th}$ percentiles and whiskers 1.5 x IQR.

### Supplementary Figure 5

RD19 expression across different co-expression samples

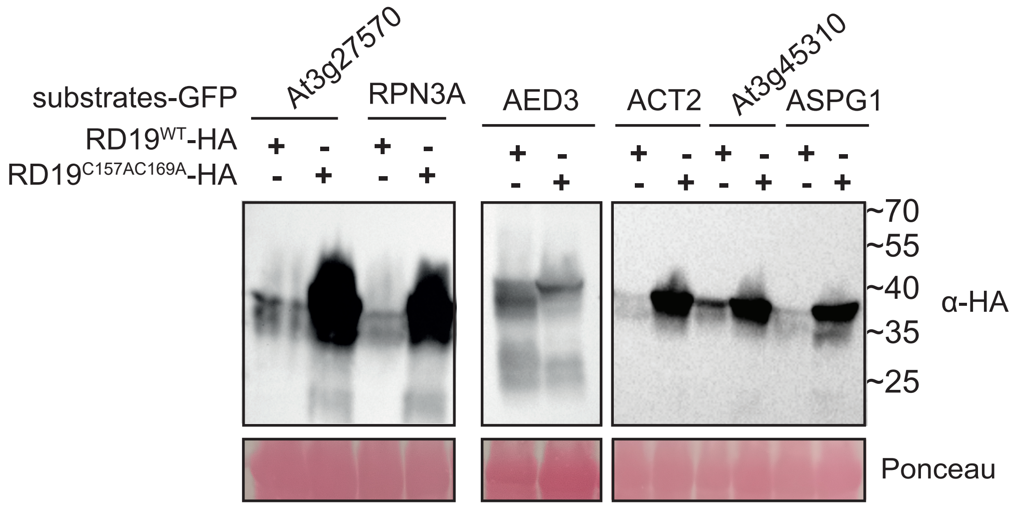


**Supplementary figure S5. RD19A protein levels across co-expression samples with candidate substrates.** Immunoblotting of co-expression of 35S::RD19A$^{WT}$-3xHA or 35S::RD19A$^{C156A_C159A}$-3xHA with substrate candidates in *N. benthamiana* detected on an immunoblot probed with α-HA antibodies. Ponceau-stained Rubisco served as a loading control, to ensure equal protein levels across samples. Images of different blots shown are representative of protein levels observed in three independent biological replicates.

### Supplementary Figure 6

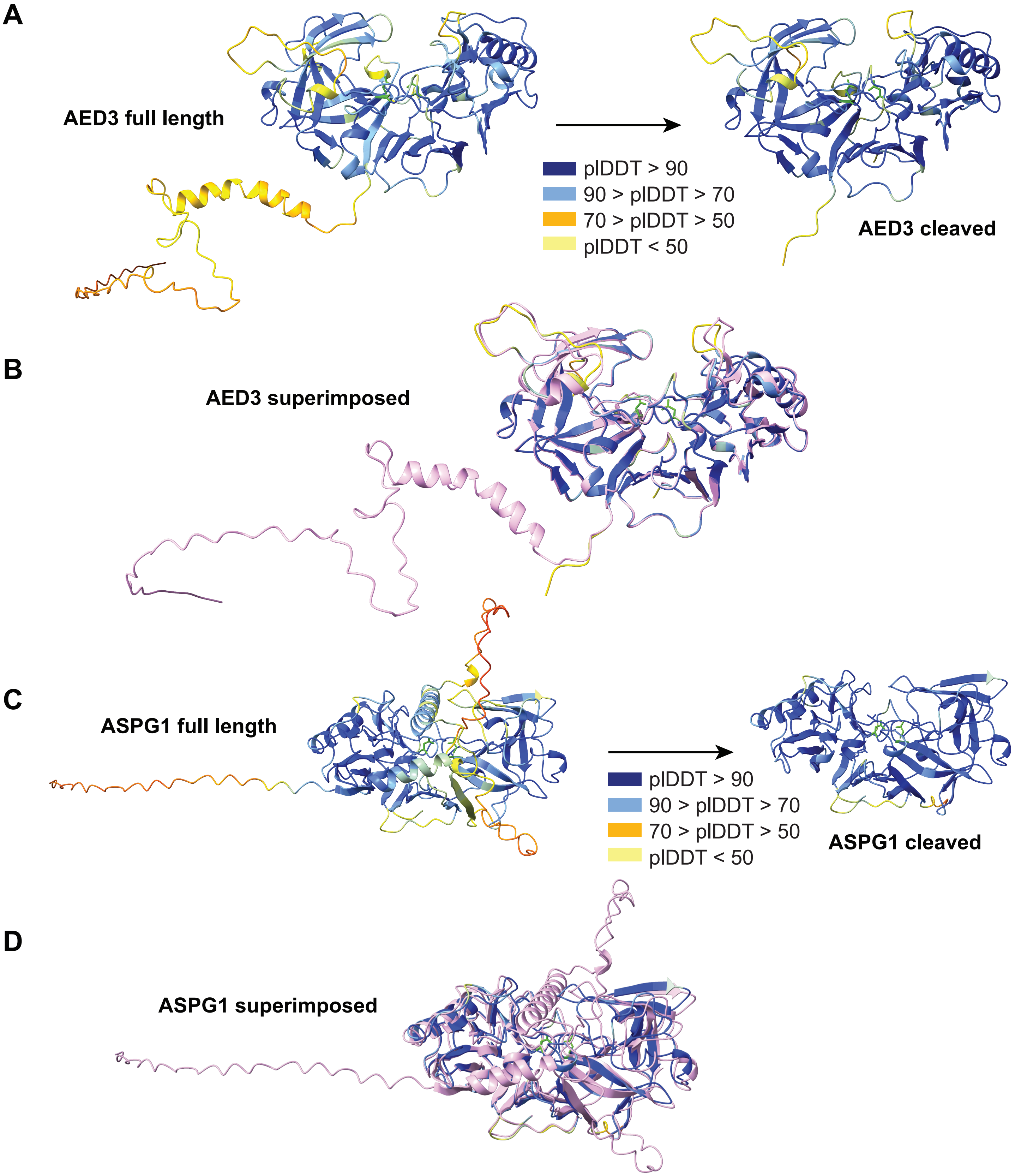


**Supplementary figure S6. AlphaFold3 modelling of AED3 and ASPG1 full length and cleaved proteins.** Structural prediction of **A)** full length (pTM = 0.92) and cleaved (pTM = 0.93) AED3 and of **B)** the two superimposed structures (RMSD score of 0.446 Å). Structural prediction of **C)** full length (pTM = 0.84) and cleaved (pTM = 0.93) ASPG1 and of **D)** the two superimposed structures (RMSD score of 1.011 Å). Structures in **A)** and **C)** are coloured based on the pIDDT score: Dark blue = Very high (pIDDT > 90), light blue = Confident (90 > pIDDT > 70), yellow = Low (70 > pIDDT > 50), orange = Very low (pIDDT < 50). In the superimposed structures of **B)** and **D)** the cleaved protein is coloured based on pIDDT scores, and full-length protein is coloured light pink.

#### Supplementary Figure 7

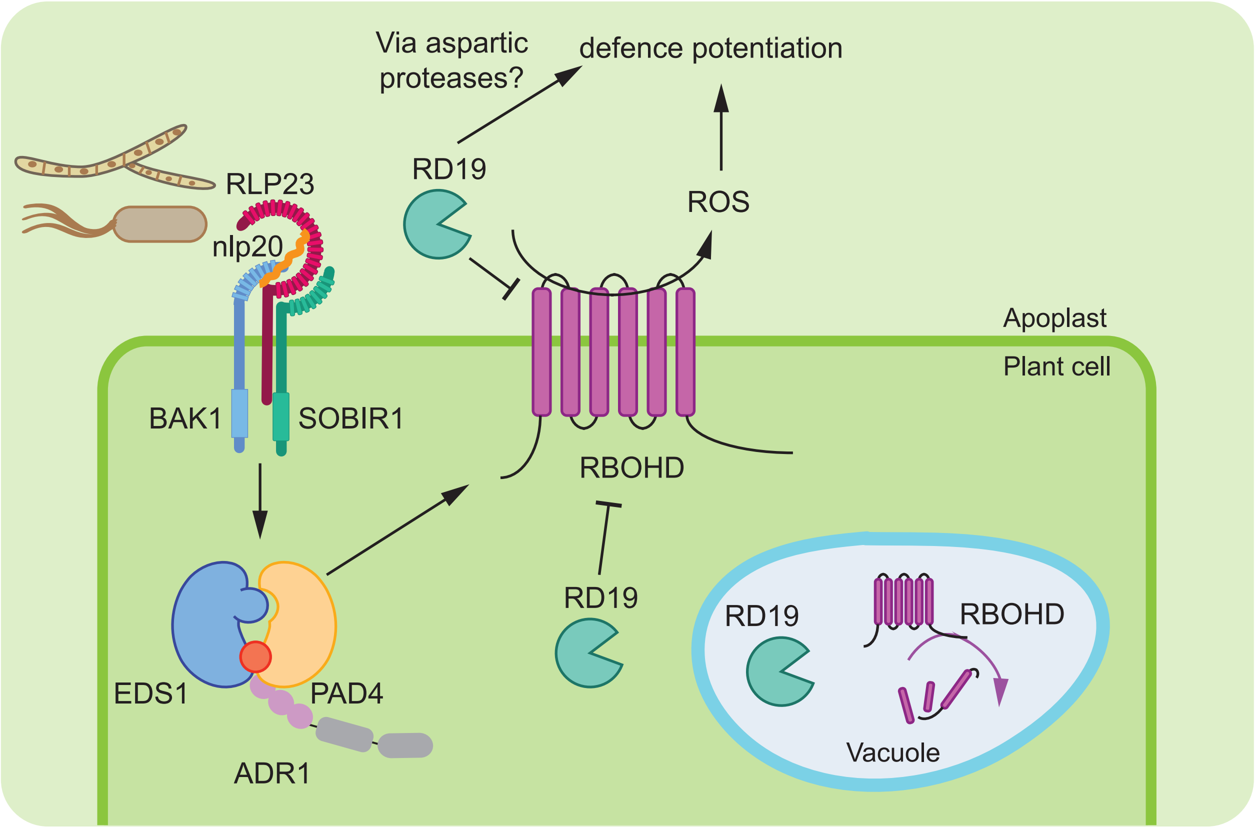


**Supplementary figure S7. Working model for RD19 activity in nlp20-triggered immunity.** Upon perception of the PAMP nlp20 by cell-surface receptor RLP23, downstream signalling via second messenger activated EDS1-PAD4-ADR1 leads to early cellular signalling outputs such as production of ROS by RBOHD at the plasma membrane. RD19 limits RBOHD protein accumulation and apoplastic ROS production, which could be through RD19 promotion of RBOHD turnover in the vacuole to prevent its recycling to the plasma membrane. Alternatively, RD19 might regulate RBOHD in the cytosol or apoplastic space. RD19 is required for nlp20-triggered defence potentiation leading to pathogen resistance, via NADPH ROS regulation and induced activation of aspartic proteases.

#### Supplementary figure legends:

**Supplementary Figure S1. Expression profile of *RD19* genes upon PTI activation and RD19A and C contributions to a PAMP-induced ROS burst**. **A)** *RD19* expression analysis of publicly available RNA-seq data sets following flg22 and nlp20 treatment. RNA-seq data sets were downloaded from the NCBI SRA data bank (flg22: PRJNA339762; GEO: GSE85923, nlp20: PRJNA549850; GEO: GSE133053). Expression of *RD19A*, *RD19B*, *RD19C* and *RD19D* are shown at h post infection (hpi) with flg22 or nlp20. Treatments were performed on 12- and 10-day old Col-0 seedlings, respectively. **B)** and **C)** Area under curve (AUC) for ROS bursts following elicitor treatments with **B)** flg22 and **C)** nlp20. Boxplots show $\log_{10}$ transformed AUC values over 60 min ROS measurements with dots of the same shape representing one biological replicate. Genotypes sharing letters above boxplots do not show statistically significant differences (ANOVA and Tukey HSD test, $p < 0.05$, n=24 (flg22 and nlp20) and n=6 (mock) leaf discs from three independent biological replicates). Centre lines of boxplots indicate the median, outer edges the $25^{th}$ and $75^{th}$ percentiles and whiskers 1.5 x IQR.

**Supplementary figure S2. Increased RBOHD protein accumulation is independent of *RBOHD* gene expression.** *RBOHD* gene expression changes were measured in non-treated (nt) leaf tissue, and at 15 min and 30 min after nlp20 infiltration. Relative quantification is represented in boxplots with dots of different shapes indicating biological replicates. Data was normalized to Col-0 mock conditions in each biological replicate. *UBI5* was used as a reference gene. Genotypes sharing letters above boxplots do not show statistically significant differences (Kruskal-Wallis and Nemenyi-Test, $p < 0.05$, n=8). Centre lines of boxplots indicate the median, outer edges the $25^{th}$ and $75^{th}$ percentiles and whiskers 1.5 x IQR.

**Supplementary figure S3. Analysis of cell-type specific expression of *RD19A* and *RD19C* from a publicly available single cell RNAseq dataset (Tang *et al.,* 2023) following *Arabidopsis* Col-0 infection with *Colletotrichum higginsianum* at 24 h and 40 h.** Mock treatment was used as a control. Heatmap depicts normalized expression values and the expression matrix was generated from monocle pipeline by aggregating expression of each gene in cells from the same cell type and normalizing this to cell number as a size factor.

**Supplementary figure S4. RD19A and C are not required for basal immune responses to *Pst* DC3000 and *Botrytis cinerea*. A)** *Pst* DC3000 growth assay in leaves of 4.5-week-old *Arabidopsis* Col-0, *pad4* and *rd19ac* at 3 d post syringe infiltration. Boxplots show colony forming units (CFU)/cm2 with each shape representing one biological replicate. Genotypes sharing letters above boxplots do not show statistically significant differences (ANOVA and Tukey HSD test, $p < 0.05$, n=24). **B)** *B. cinerea* spores were drop-inoculated on leaves of 4.5-week-old *Arabidopsis* Col-0, *rlp23* and *rd19ac* and lesion areas were measured at 2 dpi using ImageJ as a proxy of pathogen growth. Violin blots show lesion areas in $mm^2$, with dots of the same shape representing one biological replicate. Genotypes sharing letters above boxplots do not show statistically significant differences (Kruskal-Wallis and pairwise Wilcox-test, $p < 0.05$, n=90). Centre lines of boxplots indicate the median, outer edges the $25^{th}$ and $75^{th}$ percentiles and whiskers 1.5 x IQR.

**Supplementary figure S5. RD19A protein levels across co-expression samples with candidate substrates.** Immunoblotting of co-expression of 35S::RD19A$^{WT}$-3xHA or 35S::RD19A$^{C156A_C159A}$-3xHA with substrate candidates in *N. benthamiana* detected on an immunoblot probed with α-HA antibodies. Ponceau-stained Rubisco served as a loading control, to ensure equal protein levels across samples. Images of different blots shown are representative of protein levels and cleavage patterns observed in three independent biological replicates.

**Supplementary figure S6. AlphaFold3 modelling of AED3 and ASPG1 full length and cleaved proteins.** Structural prediction of **A)** full length (pTM = 0.92) and cleaved (pTM = 0.93) AED3 and of **B)** the two superimposed structures (RMSD score of 0.446 Å). Structural prediction of **C)** full length (pTM = 0.84) and cleaved (pTM = 0.93) ASPG1 and of **D)** the two superimposed structures (RMSD score of 1.011 Å). Structures in **A)** and **C)** are coloured based on the plDDT score: Dark blue = Very high (plDDT > 90), light blue = Confident (90 > plDDT > 70), yellow = Low (70 > plDDT > 50), orange = Very low (plDDT < 50). In the superimposed structures of **B)** and **D)** the cleaved protein is coloured based on plDDT scores, and full-length protein is coloured light pink.

**Supplementary figure S7. Working model for RD19 activity in nlp20-triggered immunity.** Upon perception of the PAMP nlp20 by cell-surface receptor RLP23, downstream signalling via second messenger activated EDS1-PAD4-ADR1 leads to early cellular signalling outputs such as production of ROS by RBOHD at the plasma membrane. RD19 limits RBOHD protein accumulation and apoplastic ROS production, which could be through RD19 promotion of RBOHD turnover in the vacuole to prevent its recycling to the plasma membrane. Alternatively, RD19 might regulate RBOHD in the cytosol or apoplastic space. RD19 is required for nlp20-

triggered defence potentiation leading to pathogen resistance, via NADPH ROS regulation and induced activation of aspartic proteases.

#### Supplementary table:

Table S1: Primers used in this study

| Internal ID | Sequence | Purpose | Orientation |
|---|---|---|---|
| nGH131 | GCTTCTCTAGCTAGAGTCGATCGAC | Linearize L1 vector | fw |
| nGH143 | TGTATCGATAATTGTAAATGTAATT | Linearize L1 vector | rv |
| nGH144 | CATTTACAATTATCGATACAATGGATCGTCTTAAGCTTTATTTCT | Amplification RD19 | fw |
| nGH150 | ATGGGCGGTGGTTGAGAC | Amplification RD19 | rv |
| nGH151 | TCAACCACCGCCCATTACCCATACGATGTTCCTGAC | 3xHA tag addition | fw |
| nGH152 | TCTAGCTAGAGAAGCTTATTTTCAAATTGAGGATGAGAC | 3xHA tag addition | rv |
| nGH145 | CAGGGATCTgcCGGCTCTgcCTGGAGTTTCAGCGCCACTG | SDM RD19A_C156A_C159A | fw |
| nGH146 | CAGTGGCGCTGAAACTCCAGgcAGAGCCGgcAGATCCCTG | SDM RD19A_C156A_C159A | rv |
| nGH192 | ACAATTATCGATACAATGATAACGAAATTTACATCTCCAT | Amplification At3g27570 with overhangs | fw |
| nGH193 | CTTGCTCACCATCCCACCTGACCTTCTGTAAATGCTGT | Amplification At3g27570 with overhangs | rv |
| nGH196 | ACAATTATCGATACAATGACTCAGGACGTGGAGATG | Amplification RPN3A with overhangs | fw |
| nGH197 | CTTGCTCACCATCCCAAAATCATCATCGTCTTCCTCAGCC | Amplification RPN3A with overhangs | rv |
| nGH198 | ACAATTATCGATACAATGGCCTCCTCAAGTCTCCA | Amplification AED3 with overhangs | fw |
| nGH199 | CTTGCTCACCATCCCGTTGCAGGGCTCAGGAGCA | Amplification AED3 with overhangs | rv |
| nGH202 | ACAATTATCGATACAATGGCTGAGGCTGATGATATTC | Amplification ACT2 with overhangs | fw |
| nGH203 | CTTGCTCACCATCCCGAAACATTTTCTGTGAACGATTCC | Amplification ACT2 with overhangs | rv |

| | | | |
|---|---|---|---|
| nGH206 | ACAATTATCGATACAATGTCTGTGAAACTAAACCTATCTT | Amplification At3g45310 with overhangs | fw |
| nGH207 | CTTGCTCACCATCCCGGCTACAACCGGATACGATGA | Amplification At3g45310 with overhangs | rv |
| nGH204 | ACAATTATCGATACAATGGCTTTCCCGCGATTTC | Amplification ASPG1 with overhangs | fw |
| nGH205 | CTTGCTCACCATCCCGCATTTATTCCCTGACAATCCG | Amplification ASPG1 with overhangs | rv |
| nFL085 | CGAATGGCATCCTTTCTCAATC | qPCR_RBOHD | fw |
| nFL086 | GTCACCGAGAGTGCGGATATG | qPCR_RBOHD | rv |
| nFL170 | TCGCTGGCATAACACTATGG | qPCR_PAD3 | fw |
| nFL171 | TTGGGAGCAAGAGTGGAGT | qPCR_PAD3 | rv |
| nFL005 | TTAAACTCGACGATGCAACA | qPCR_FRK1 | fw |
| nFL006 | GATGGAAGTTTTCCCGTTTT | qPCR_FRK1 | rv |

#### Supplementary methods:

**Proteomic Identification of Protease Cleavage Sites (PICS)** *E. coli* proteome derived peptide libraries were obtained by digestion of *E. coli* total cell lysates with either Trypsin or GluC. Three library aliquots with 50 µg concentration each were incubated with apoplastic fluids from *N. benthamiana* expressing proteolytic active RD19A, or a catalytically inactive RD19A-mutant or RD19A pre-treated with 100 µM E64 (in 10 mM $NaPO_4$, 150 mM NaCl, 1 mM EDTA, 0.5 mM DTT, 50 µM PepstatinA (Roth) and 50 µM 3,4-Dichlor-isocumarin (Sigma)) at room temperature for 1 h. The reaction was stopped by addition of 100 µM E64 and freezing samples in liquid nitrogen. Peptides were then differentially stable isotope labelled with different combinations of formaldehyde and cyanoborohydride isotope ($CH_2O$ and $NaBH_3CN$ for inhibited RD19A; $CD_2O$ and $NaBH_3CN$ for active RD19; $^{13}CD_2O$ and $NaBD_3CN$ for inactive mutant RD19A, each 30mM). Labelling reactions were quenched with Tris-HCl pH 6.8, combined and subsequently used for mass spectrometry data acquisition and analysis.

**High-efficiency Undecanal-based N-Termini EnRichment (HUNTER).** For each sample, protein concentration was determined using the Bicinchoninic Acid (BCA) Protein Assay (Thermo). For each replicate and condition, 200 µg proteins were reduced, alkylated, and free primary amines at protein n-termini and Lys side chains dimethylated with heavy formaldehyde ($^{13}CD_2O$) and cyanoborohydride ('single-heavy label'). The reaction was quenched by adding 100mM ammonium bicarbonate and stored at 37°C for 30 min. Proteins were digested with Trypsin in a 1:100 protease:protein ratio over night at 37°C. Primary amines at trypsin-generated peptide N-termini were modified with undecanal and removed by solid phase extraction as described (Demir *et al.*, 2022*b*), leaving N-terminal peptides enriched in the flow through. All peptide samples were acidified and desalted by using STAGE tips (Rappsilber *et* *al.*, 2007) self-packed with SDB-RPS material (Empore, 3M) and subsequently analysed by mass spectrometry.

**Mass spectrometry data acquisition.** PICS samples were measured with an Orbitrap Elite mass spectrometer instrument coupled to an UltiMate 3000 RSLCnano HPLC system (Thermo). Peptides were separated on a two-column setup with a C18 trap (nanoEase M/Z Symmetry, 20 mm x 180 µm inner diameter, Waters) and analytical columns (nanoEase M/Z HSS C18 T3, 250mm x 75 µm inner diameter, Waters) at a flow rate of 0.3 µl/min and column temperature of 40°C. Peptide elution was controlled by applying a binary solvent gradient. Solvent A consisted of 0.1% (v/v) formic acid (FA), while solvent B consisted of 0.1% (v/v) FA, 50% (v/v) methanol (MeOH), and 30% (v/v) acetonitrile (ACN). The elution gradient went from 7% to 55% solvent B over 175 mi, followed by a 55% to 95% increase in 35 min, and finally held at 95% for 5 min. This system was interfaced online with the Nanospray Flex ion source featuring the PST-HV-NFU liquid junction (MS Wil) and fused silica emitters (EM-20-360; MicrOmics Technologies LLC). The following parameters were applied for MS1 scans: mass range of m/z 370 - 1,700, a resolution of 120,000 (at m/z 400), a target value (AGC) of 1 x $10^6$ ions, and a maximum injection time (IT) of 200 ms. For MS2 scans, up to 15 most intense precursor peptides with a charge ≥ 2 were selected for collision induced dissociation (CID) in the linear trap with a normalized collision energy of 35%, activation time 10 sec q-value = 0.25, an AGC of 5000 ions, a max. IT of 150 ms, and dynamic exclusion for 45 s.

HUNTER samples were measured with an Exploris 480 mass spectrometer coupled to an UltiMate 3000 RSLCnano HPLC system (Thermo). Peptides were separated on a two-column setup with a µPAC C18 trapping column and a µPAC pillar array analytical column of 50 cm length (Thermo) at a flow rate of 0.5 µl/min and column oven temperature of 40°C. Peptide elution was controlled using a binary buffer system consisting of A (0.1% formic acid) and B (86% acetonitrile, 0.1% formic acid). The elution gradient went from 1% to 4% solvent B over 2 min, 4% to 25% solvent B over 24 min, followed by a 25% to 44% increase in 12 min, and a final flush held at 90% for 4 min. This system was interfaced online with the Nanospray Flex ion source (Thermo) featuring the PST-HV-NFU liquid junction (MS Wil) and fused silica emitters (EM-20-360; MicrOmics Technologies LLC). MS data was acquired in data independent acquisition (DIA) mode with one overview spectrum (RF lens of 40%, normalized AGC target of 300%, maximum injection time of 45 ms, m/z range of 350 to 1.400, resolution of 120.000, profile mode) and MS2 fragment spectra with 14 Da isolation windows at a precursor mass range of 361-450 m/z, 7 Da windows in a mass range of 450-800 m/z and 14 Da windows in a mass range of 800-1100 m/z each with a window overlap of 1 m/z and a resolution of 30000 and a maximum injection time (IT) of 54 ms.

**Mass spectrometry data analysis**

For HUNTER data, peptide sequences were identified with FragPipe (v. 22; Yu et al., 2023) essentially as described (Stockert *et al.*, 2024). Enzyme specificity set to "semi-specific" ArgC. N-terminal dimethylation (+34.0631) and Carbamidomethylation of Cys residues (+57.02146) were set as fixed modifications, N-terminal acetylation (+42.0106), N-terminal pyroGlu formation either from glutamic acid (-18.0106) or glutamine (-17.0265), methionine oxidation

(+15.9949) and N-terminal loss of ammonia from cysteine (-17.0265) were set as variable modifications. LFQ and “Match between runs” was enabled. Spectra were matched against an UniProt reference proteome for *Arabidopsis thaliana* (accessed October 10, 2023). One sample of the wildtype untreated condition was excluded from data analysis due to measurement error.
